# Emergent perceptual biases from state-space geometry in spiking recurrent neural networks trained to discriminate time intervals

**DOI:** 10.1101/2022.11.26.518023

**Authors:** Luis Serrano-Fernández, Manuel Beirán, Néstor Parga

## Abstract

A stimulus held in working memory is perceived as contracted towards the average stimulus. This contraction bias has been extensively studied in psychophysics, but little is known about its origin from neural activity. By training recurrent networks of spiking neurons to discriminate temporal intervals, we explored the causes of this bias and how behavior relates to population firing activity. We found that the trained networks exhibited animal-like behavior. Various geometric features of neural trajectories in state space encoded warped representations of the durations of the first interval modulated by sensory history. Formulating a novel normative model, we showed that these representations conveyed a Bayesian estimate of the interval durations, thus relating activity and behavior. Importantly, our findings demonstrate that Bayesian computations already occur during the sensory phase of the first stimulus and persist throughout its maintenance in working memory, until the time of stimulus comparison.

## Introduction

Perception is not based purely on the current stimulus but is affected by other factors such as recent stimulation, prior knowledge and context, giving rise to perceptual biases [1]–[9]. Examples of these biases appear in magnitude estimation [10]–[13] and delayed comparison tasks [14]–[17], in which a stimulus parameter is perceived as shifted toward the mean of its distribution. These phenomena, known respectively as the central tendency and the contraction bias, have been extensively investigated in psychophysics [10]–[17] and, more recently and to a lesser extent, with neural activity recordings in temporal reproduction tasks [18]–[24] and delayed comparison tasks [6], [9], [25]. However their causes have not been fully elucidated, particularly in the latter.

The contraction bias is affected by the sensory history [6], [26], [27]. A remarkable study found that in delayed comparison tasks the activity of posterior parietal cortex (PPC) neurons carries information about past stimuli, but not about the current stimulus [6]. Furthermore, that activity was causally related to the contraction bias.

Within a Bayesian framework, prior experience is integrated with the noisy observation of the current stimulus. This integration would result in a sensory representation biased to the most reliable or likely stimuli, generating the contraction bias in delayed comparison tasks [9], [15], [26], [28], [29]. However, despite their relevance, no study has addressed the fundamental questions of how the integration of those relevant factors takes place in neural circuits and what features of neural population activity relate to behavior. Moreover, it is not known how the neural information about the preceding stimuli relates to Bayesian computations.

New computational techniques enable us to train recurrent neural networks on tasks similar to those used in the experiments [4], [30]–[35]. This approach provides us with a valuable tool to investigate the network mechanisms responsible for perceptual biases. Based on trained networks, we can formulate hypotheses and draw predictions to guide novel experiments. In addition, trained networks allow full access to the state-space dynamics and geometry of the neural trajectories and thus to the study of the relationship between population activity and behavior [20], [21].

In this work, to further the understanding of the causes of the contraction bias and its connection with neural population activity, we trained spiking recurrent neural networks (sRNNs) with the full-FORCE algorithm [36], [37](see also [38]) to compare the durations of two temporal intervals (*d*_1_ and *d*_2_), spaced by a delay interval. The trained networks did exhibit the bias. Examination of the networks’ behavior allowed us to identify potential sources of the contraction bias: short- and long-term sensory history and the integration of noisy observations with prior knowledge. Accordingly, an ideal observer would have access to all of them; this led us to propose a novel Bayesian normative model that can be used to guide the analysis of the data and help identify the network computations responsible for the emergence of the contraction bias.

The activity of the neural population described trajectories in a low-dimensional state space. The geometry of the trajectories during the task encoded the durations of the first interval but did so by compressing them with respect to their true values. These compressions were hypothesized to convey a Bayesian estimate of the interval, thus relating population activity and behavior. Indeed, the Bayesian estimator of *d*_1_ showed similar compressions as those of the state-space trajectories, thereby supporting a Bayesian explanation of the contraction bias.

Our work leads to testable predictions about the causes of the contraction bias and how they are interrelated. It predicts a relationship between geometric properties of state-space trajectories and behavior. Moreover, the normative model predicts that the sensory history contributes not only to the mean of the observations of the first interval but also modulates their variance; this modulation has behavioral and physiological consequences, since an increase in observation variance implies a larger contraction bias and affects the geometric properties of the trajectories. These findings are a consequence of the recurrence of the network and the existence of a delay period between the two presented stimuli. More generally, then, we expect the contraction bias be explained in a similar way in other two-interval forced-choice tasks.

Our research significantly contributes to the understanding of perception by examining the impact of current and recent stimulation, prior knowledge, and contextual information, which collectively give rise to perceptual biases. Notably, our findings highlight the predominant role of Bayesian principles in shaping perception. However, we found that, contrary to conventional assumptions, the integration of prior information with the current noisy stimulation is not confined solely to the short-term memory stage but is already active during the initial sensory processing phase. We observed its presence across various geometric features of population activity, from the moment of stimulation until the comparison with a second stimulus is made. Furthermore, our work sheds light on the influence of recent sensory history or noisy perturbations and their impact on contraction bias, which manifest as modulations of Bayesian processing.

## Results

### Network training and numerical experiments

We trained sRNNs to decide which of two time intervals, *d*_1_ and *d*_2_, presented sequentially and separated by a delay interval, was longer (Fig. 1a). Networks were trained with a set of duration pairs (*d*_1_, *d*_2_), also named classes (Fig. 1b), selected randomly and independently in each trial, using the full-FORCE algorithm for sRNNs [36]–[38] (Supplementary Fig. S1). Trials were presented continuously, without altering the network activity at the end of each trial. Once a network was trained, a large number of test trials were obtained for further analyses of task performance and population activity (Methods). We considered three different trial blocks: a block in which only diagonal classes (classes labeled from 1 to 10 in Fig. 1b) were tested, a block where only horizontal classes (*d*_2_=370 ms) were tested and a block where all classes were tested (Methods). These will be referred to as the diagonal, horizontal and full blocks (Fig. 1b). Each block defined a context with a distinct prior distribution of *d*_1_ (Fig. 1c); note that some classes appeared in more than one context.

**Fig. 1.**
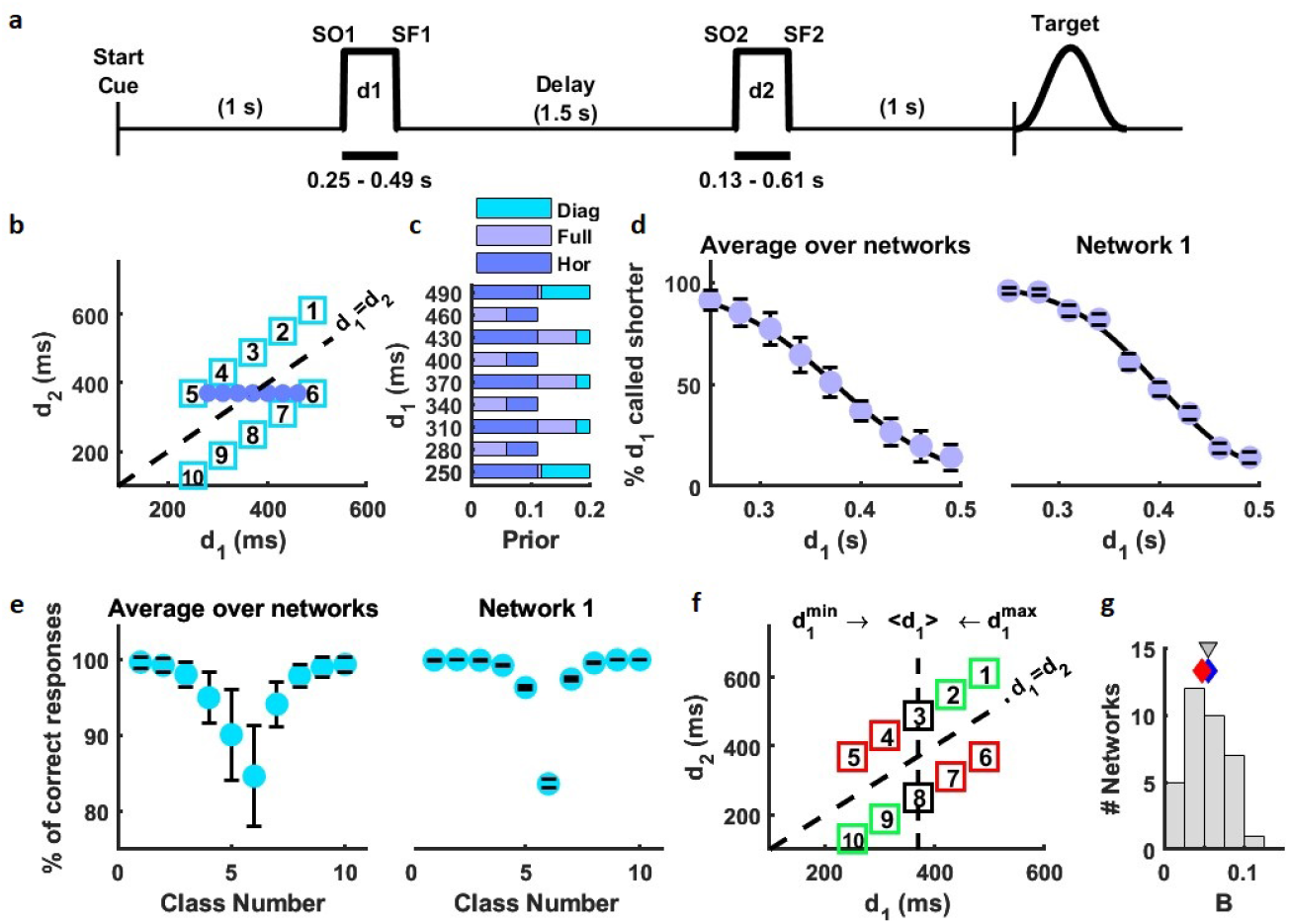
Task and network behavior. **a**. Timeline of the time interval discrimination task. The Start Cue indicates the beginning of the trial. One second later, two time intervals (*d*_1_ and *d*_2_) are presented sequentially, separated by a delay period of 1,500 ms (SO1 and SO2 are their respective onset times, while SF1 and SF2 denote their offset times). During the training epoch, a target (a beta distribution) is presented one second after SF2. **b**. Stimulus set of possible classes or pairs (*d*_1_, *d*_2_), indicated by squares or blue circles. There are three block types: horizontal block (all classes with *d*_2_=370 ms), diagonal block (classes labeled from 1 to 10) and full block, defined by all the classes. Classes 5 and 6 are common to the three blocks. **c**. Prior distributions of *d*_1_ in each block. **d**. Psychometric curves, computed using test trials with horizontal classes from the full block. **Left**: Averaged psychometric curve over 35 networks trained and tested on consecutive trials without activity reset. Error bars are standard deviations over networks. **Right**: Psychometric curve for example network 1. Error bars were computed with 1,000 bootstrap resamples. Black lines in both psychometric curves are sigmoidal fits (Methods). **e**. Percentage of correct test trials, computed on the diagonal block. **Left**: Average accuracy over 35 networks. Error bars indicate standard deviations computed over networks. **Right**: Accuracy for example network 1. Error bars were computed with 1,000 bootstrap resamples. **f**. Contraction bias: the first interval (*d*_1_) is coded contracted towards the mean of its distribution. This bias induces errors in some classes (red squares) while favors others (green squares). **g**. Histogram over the networks of the phenomenological bias (*B*) (Methods, Eq. (14)). Gray triangle indicates the histogram mean. Diamonds indicate *B* for example network 1 (blue) and example network 2 (red).

In the main numerical experiment we trained a total of 35 networks (obtained from random initializations of the network parameters), with a fixed delay interval of 1,500 ms. Unless otherwise specified, the analyses were performed on this experiment. In a separate numerical experiment, to examine the effect of the delay period duration on the contraction bias, we trained 15 sRNNs by selecting its duration on each trial randomly from a set of four possible values, respectively (Methods). In another experiment, we trained 7 networks resetting the population activity at the end of every trial (Methods).

### The trained networks exhibited the contraction bias

The psychometric curves were computed using test trials from the horizontal classes in the full block. Fig. 1d shows this curve for their mean over all trained networks (left) and for example network 1 (right) (see Extended Data Fig. 1a for example network 2). Psychometric curves were heterogeneous across the 35 trained networks (Supplementary Fig. S2). Next we analyzed the discrimination performance using test trials from the diagonal block. Note that classes in this block have a fixed duration difference Δ*d* = |*d*_1_ − *d*_2_| (60 ms; Fig. 1b). However, contrary to what would be expected for classes with the same duration differences, accuracy was not uniform, as seen in its average over networks (Fig. 1e, left) and in the two example networks, Fig. 1e (right) and Extended Data Fig. 1b, respectively. The contraction bias can explain this disparity. This bias is usually described as a shift of the encoding of duration of the first stimulus towards the mean of its distribution (i.e., ⟨*d*_1_⟩=370 ms, Fig. 1f). This means that a short duration (i.e., *d*_1_=250 ms) is encoded as longer, while a long duration is encoded as shorter. Therefore, correct decisions for classes with *d*_1_ *< d*_2_ are increasingly impaired by the bias as *d*_1_ decreases (upper diagonal, Fig. 1f), while classes with *d*_1_ *> d*_2_ are increasingly facilitated by the bias (lower diagonal, Fig. 1f). To make this bias structure explicit, we numbered the classes such that those with favorable bias effects are near the extremes, while classes with disfavorable effects are around the center (Fig. 1f). With this arrangement of classes the decision accuracy was higher for classes near the two extremes (Fig. 1e). Note that since the networks were trained with sequences of temporally uncorrelated *d*_1_ values, this result implies that correlations in the training stimuli are not crucial for generating the bias.

The contraction bias of a network was quantified as the root-mean-square deviation of the performance, relative to the diagonal classes with *d*_1_ = 370 ms, a quantity denoted as *B*(Methods). The distribution of this bias metric spanned approximately one order of magnitude across networks (Fig. 1g).

### Behavior and possible causes of the contraction bias

We postulate three different sources for the contraction bias. One possible explanation is inter-trial effects, through the persistence of the neural representations of stimuli applied in previous trials. This would produce a contraction driven by the mean of the stimulus parameter or by the most recently presented stimuli [6], [26]. In addition, it has been proposed that the bias can be generated by a Bayesian mechanism that occurs if, at the time of comparison of the two stimuli, the uncertainty in the observation of the first stimulus is greater than the uncertainty in the observation of the second one -which creates a tendency to rely more on the *a priori* knowledge on the first stimulus [15]. In support of this idea, behavioral data from monkeys discriminating tactile frequencies were well fit with a Bayesian model [9]. Finally, the stimulus statistics could be stored in the network synapses during the learning stage, then affecting behavior with a contraction of the intervals toward the learned mean [28]. We describe below the results of numerical experiments performed to examine these three possible causes of contraction bias.

To check that the class in the previous trial can modulate behavior, we sorted a large number of test trials by the preceding class and calculated the choice accuracy for each case (Methods). The analysis confirmed that the preceding trial modulates performance (see Fig. 2a and Extended Data Fig. 1c for example networks 1 and 2, respectively). In addition, we observed the presence of a choice bias, which depended on whether the preceding choice had been rewarded (Methods).

**Fig. 2.**
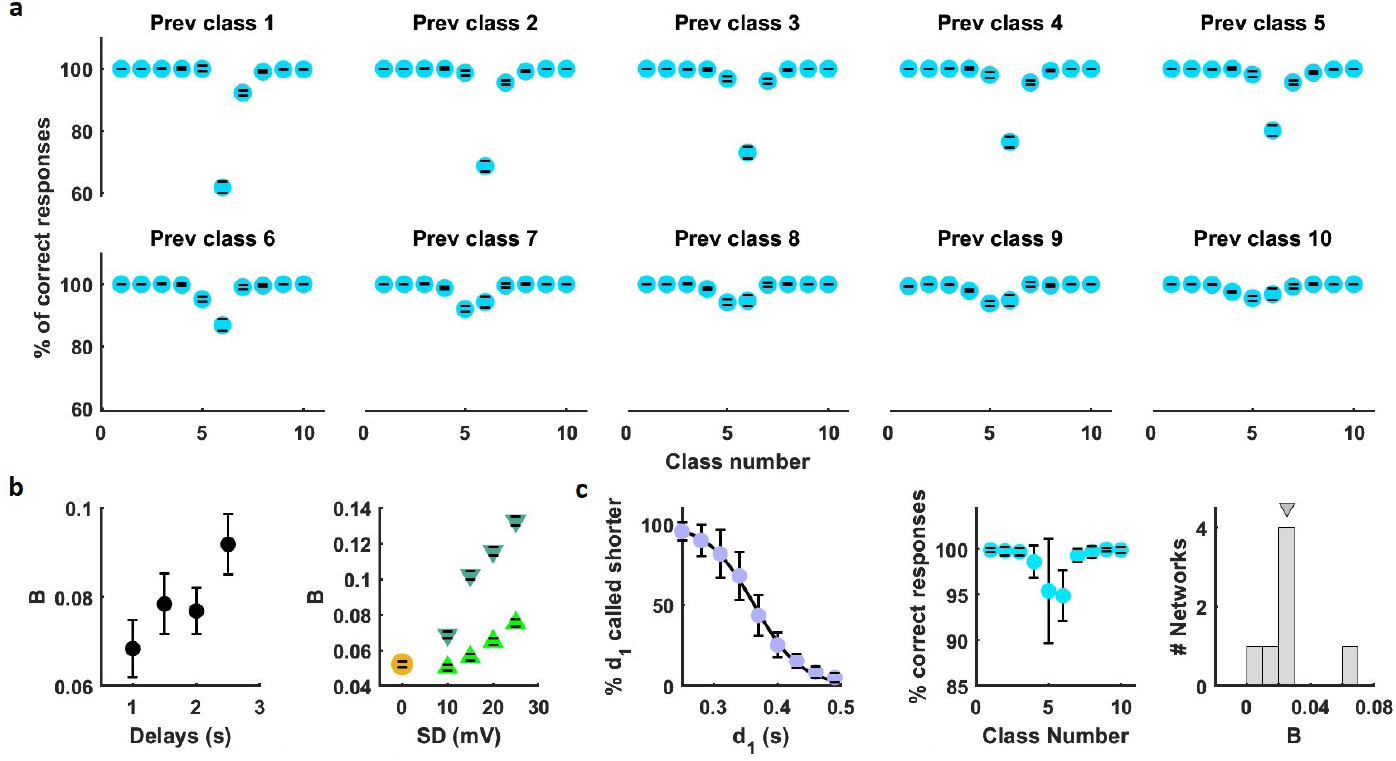
Potential causes of the contraction bias. **a**. Accuracy depends on the preceding class (main numerical experiment). Accuracy conditioned on each of the ten possible previous classes (example network 1; test trials in the diagonal block). Error bars computed with 1,000 bootstrap resamples. **b. Left**. Phenomenological bias (*B*) increases with delay-period duration (second numerical experiment). *B* vs. delay-period duration for a network trained by randomly interleaving four different delay durations (1,000, 1,500, 2,000 and 2,500 ms) and tested in four blocks of test trials, each one with a fixed delay duration (diagonal block). Error bars computed with 100 bootstrap resamples. **Right**. *B* versus the standard deviation of applied Gaussian noise. Contraction bias increases with Gaussian noise applied to the membrane potential of LIF neurons from example network 1. Upward-pointing light green triangles refer to the noise applied during the presentation of the first stimulus while downward-pointing dark green triangles refer to the noise applied during the delay period. Orange circle indicates the case without noise. Error bars computed with 1,000 bootstrap resamples. **c**. Contraction bias in the absence of sensory history, such that the neural activity was reset to a different random initial state in all training and test trials (Methods). **Left**: Average over 7 networks of the psychometric curves. Horizontal classes collected from the full block. Black line is a sigmoidal fit (see Extended Data Fig. 2a for a single network). **Middle**: Average over 7 networks of the accuracy. Classes from the diagonal block. Error bars are standard deviations over networks (see Extended Data Fig. 2b for an example network). **Right**: Histogram of the bias *B*(Methods, Eq. (14)), from all 7 reset networks. Gray triangle indicates the histogram’s mean.

If the contraction bias were due solely to the preceding stimulation then it should remain constant or weaken as the duration of the delay period is increased [29]. However, if a Bayesian mechanism were operating in our networks, we would expect that as the duration of the delay epoch increases, the noise affecting the representation of the first interval would also increase, resulting in a larger contraction bias [29]. To compare the bias for different durations of the delay period, we trained 15 networks to perform time interval discrimination for four delay durations, randomly interleaving these values during training (Methods). After training, blocks of test trials (diagonal classes) with fixed delay period duration were collected and analyzed. We found that the bias *B* increased with the delay duration (Fig. 2b, left). Additionally, it is expected that the contraction bias would increase when noise is introduced into the trained networks. To investigate this, we perturbed network 1 with external noise selectively either during stimulus presentation or during the maintenance of the stimulus in working memory (Methods). Consistent with Bayesian concepts, we observed an increase in B in both scenarios (Fig. 2b, right). This finding suggests that Bayesian processing already occurs during the sensory phase of the task.

To gain further insight into the mechanisms that could account for the contraction bias we trained networks by resetting the network activity at the end of each trial, rather than maintaining continuity of activity during the inter-trial epoch (Methods). If the bias only depended on the information about past trials, then the bias would disappear in the absence of this information. In case the bias persisted, then we would have to look for some other mechanism capable of producing it. To answer this question we investigated the seven networks trained under this condition. The accuracy of six of them showed a clear bias, as can be observed in the network mean of the psychometric function (Fig. 2c, left) and the accuracy (Fig. 2c, middle). It is noteworthy that the bias *B* turns out to be weaker than in the networks trained with no reset (Fig. 2c, right).

So far we have analyzed behavioral data from trained networks with an eye to potential sources of the contraction bias: sensory history, noisy observations and prior knowledge and memorized stationary statistics. We will return to these results later (section “Ideal observer model”), to examine them in light of a normative model that incorporates the presumed causes of contraction bias. Before that we examine whether these factors were present in the neuronal activity.

### State-space dynamics and geometry

We adopted the view that neural circuits represent and process information through the state-space dynamics of neural populations [4], [30], [32], [33], [39]– [43]. Most analyses on neural firing activity were performed at the population level, the population being all those neurons encoding *d*_1_ (Methods). We projected the population activity (z-score) onto the directions in state space responsible for at least 90% of the data variance (Methods). As expected, during the presentation of the first interval the population z-score followed a common path until the end of the duration of the respective *d*_1_, defining a curved manifold (Fig. 3a). This feature also appeared in recordings of monkeys performing time reproduction tasks, during the presentation of the sample interval [19]–[21]. The similarity is not surprising, since both the first interval in the discrimination task and the sample interval in the reproduction task must be represented in the network activity to complete the corresponding task. The trajectories evolved at different speeds during the delay period, so that the longer the duration of their interval *d*_1_, the higher the speed (Fig. 3b). As the trajectories approached the end of the interval their speeds decreased (Fig. 3c), reaching *d*_1_-dependent terminal states that appear in orderly alignment (Fig. 3b, inset). Correspondingly, the percentage of variance explained by the first principal component approached 100% (Fig. 3c, inset), showing that the trajectory arrived to a one-dimensional quasi-attractor region. However, this result does not specify which estimator of *d*_1_ is maintained by the network when it approaches the quasi-attractor. Addressing this question, as well as what geometrical properties of the trajectories encode the estimator, requires connecting behavior with population activity and will be dealt with later (section “State-space geometry is related to behavior”).

**Fig. 3.**
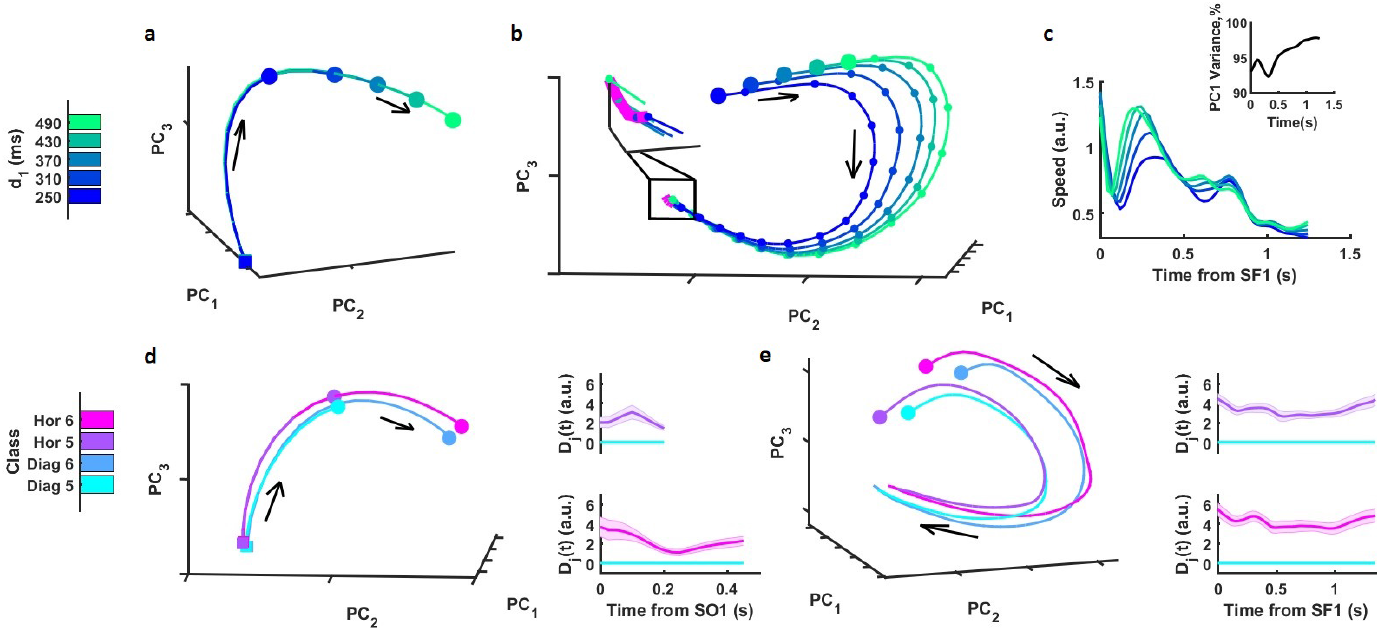
State-space dynamics and contexts. **a-e**. Example network 1. **a-c**. Projection of neural trajectories onto the first three PCs. **a**. Curved manifold during presentation of *d*_1_. Squares indicate SO1 and circles indicate SF1. **b**. Neural trajectories during the delay period. Big circles indicate the SF1s. Small circles indicate neural states at 100 ms increments. Note that the distance between consecutive circles reflects speed. **Inset** shows the quasi-attractor (magenta line), represented by connecting the terminal states at the end of the delay period. **c**. Speed of neural trajectories versus time, during the delay period (Methods). **Inset** shows the percentage of variance explained by the first PC. By the end of the delay period its variance approached 100%. **d-e**. Comparison of diagonal and horizontal contexts using the classes 5 (*d*_1_=250 ms) and 6 (*d*_1_=490 ms), which are common to both blocks. **d**. Geometry during the presentation of the first interval. **Left**: State-space trajectories of classes 5 and 6. **Right**: Relative distances of the classes 5 (upper) and 6 (bottom). The number of PCs needed to explain at least 90% of variance was two for class 5 and three for class 6. Shadings indicate the median ± 95% CIs, computed from 100 bootstrap resamples. **e**. Geometry during the delay period. **Left**: State-space trajectories of classes 5 and 6. **Right**: Relative distances of classes 5 (upper) and 6 (bottom). Four PCs were needed to explain at least 90% of variance. Shadings are as in (d), right. All the analyses were done with a time window of 150 ms sliding every 25 ms.

If the networks were to implement a Bayesian computation, integrating the noisy measure of the current *d*_1_ with prior knowledge, then, when the same value of *d*_1_ occurs in two different contexts, the population activity should follow distinct trajectories. Since classes 5 and 6 are shared by the horizontal and diagonal blocks (Fig. 1b), we wondered whether the networks learnt that these blocks define different contexts (*h* and *d*). To verify this supposition we first defined relative distances following the Kinematic analysis of Neural Trajectory (KiNeT) methodology [19]. Briefly, for each class shared by the blocks *h* and *d*(that we indicate by *j* = 5 or 6) we considered its trajectories in the two contexts (trajectories *j*_*h*_ and *j*_*d*_). Then we selected one of them (e.g., *j*_*d*_) as a reference and defined corresponding states between the reference and the other trajectory (*j*_*h*_). If at time *t* after the input pulse (SO1 or SF1, see Fig. 1a) the reference trajectory was in the state *s*_*ref*_ (*t*), the corresponding state in trajectory *j*_*h*_ was taken as the state *s*_*j*_ closest to *s*_*ref*_ (*t*). Then, for each time *t*, we computed the Euclidean distance *D*_*j*_ (*t*) between the pair of corresponding states. During the presentation of the first time interval, trajectories of classes shared by the two contexts (Fig. 3d, left. Example network 1) evolved disjointly, keeping a significantly non-zero relative distance from each other (Fig. 3d, right). During the delay period, those trajectories (Fig. 3e, left) also maintained significantly non-zero distances (Fig. 3e, right). These results indicate that the networks acquired knowledge about the context during the training stage. It should be noted that the networks were not trained specifically on the horizontal and diagonal contexts, since all the classes of the set in Fig. 1b, chosen randomly, were used for that purpose (Methods).

### Neural activity is affected by sensory history

The results in Figs. 3d,e point to the influence of sensory history on the current trial. We now study this question in a systematic and quantitative way. We first checked whether the neural activity in the current trial maintained information about the interval *d*_1_ presented in the preceding ones. To do so, we began with an analysis of the trajectories in state space, and then performed a more quantitative analysis of mutual information. Note that if the neural population coded the value of the current *d*_1_ throughout the delay period, then trajectories corresponding to different durations should stay at significantly non-zero relative distances. Likewise, if the representation of the first interval persisted to the next trial, trajectories should still maintain non-zero distances during the inter-trial-interval (ITI). Again, we employed the KiNeT methodology [19]; for each pair of trajectories (*j* and *j* + 1) we selected trajectory *j* + 1 as a reference and, for each time *t*, we computed the Euclidean distance *D*_*j*_ (*t*) between the pair of corresponding states (Methods). All pairs of consecutive trajectories in the diagonal set maintained non-zero relative distances during the delay period (Fig. 4a) and ITI (Fig. 4b). This implies that the neural representation of the first interval is maintained beyond the epoch in which it is presented.

**Fig. 4.**
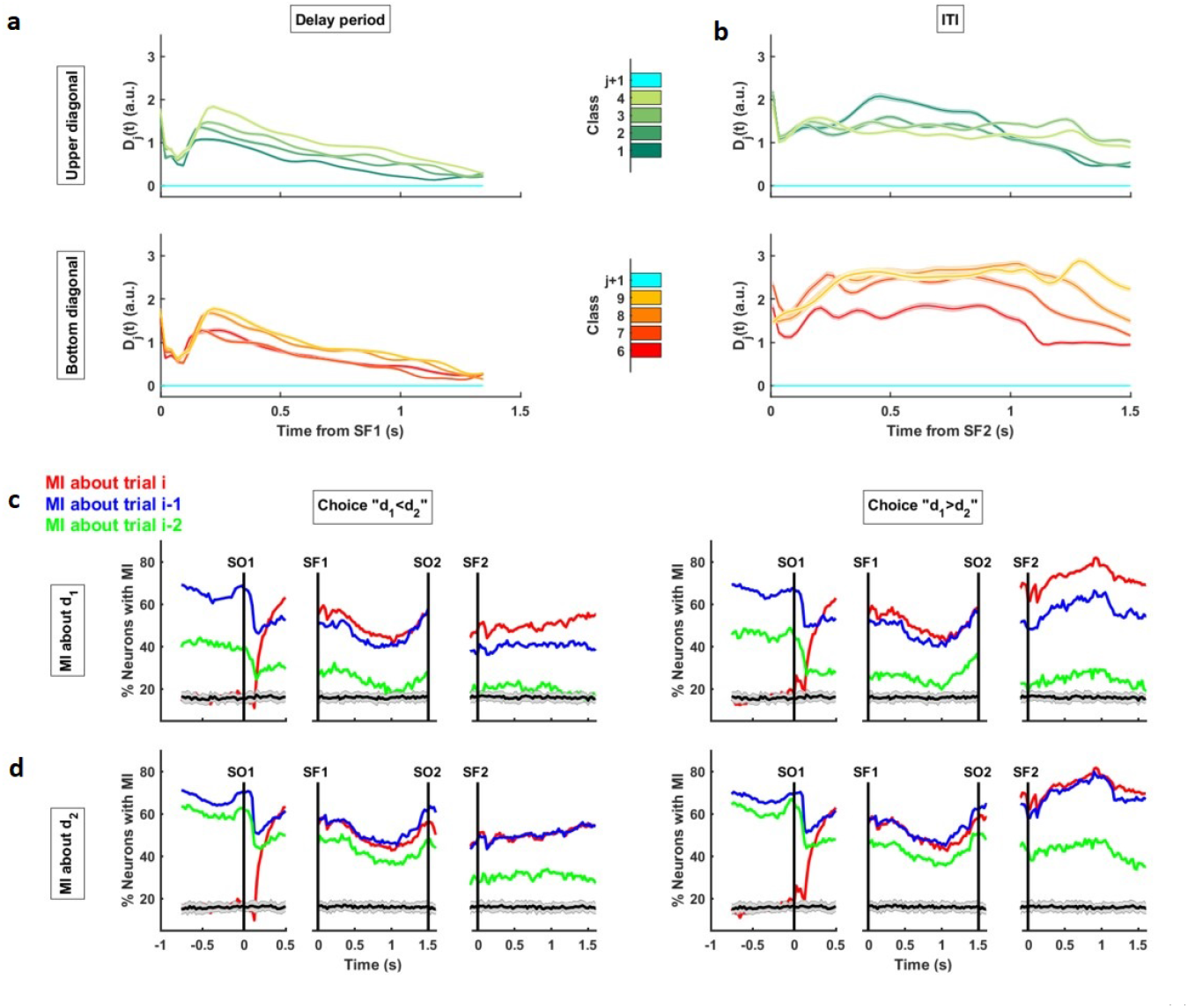
Mutual Information analyses. **a-d** Example network 1. **a-b**. Relative distances *D*_*j*_(*t*) between pairs of consecutive classes from diagonal block (*j* + 1 denotes the reference class) during the delay period (a) and the ITI (b). The (rather thin) shadings indicate the mean ± 95% CIs computed from 100 bootstrap resamples. **c-d**. Percentage of neurons with significant mutual information about the first (c) and second (d) time interval (red: from current trial *i*; blue: from trial *i* − 1; green: from trial *i* − 2), conditioning on correct trials and on choice (**left**: choice “*d*_1_ *< d*_2_”; **right**: choice “*d*_1_ *> d*_2_”). Horizontal gray shadings refer to the mean and standard deviation computed from shuffled data (Methods).

To further support this conclusion we analyzed the mutual information (MI) that the activity had on the presented time intervals. Most neurons maintained significant information about the current *d*_1_ during the delay period and even during the following ITI (Supplementary Fig. S3). During the current trial there was a fraction of neurons showing significant MI about the first interval in the current (Fig. 4c, red lines), the preceding (blue lines) and the second previous trial (green lines). Soon after SO1 (after a time equivalent to the shortest first interval), the fraction of neurons with MI about the current *d*_1_ increased rapidly and remained high during both the entire delay period and the ITI. The population of neurons with significant MI on the first interval of the preceding trial (blue), was quite large before SO1, during the delay period it stayed at a similar level as the population with MI on the current *d*_1_, although it became smaller during the ITI. In contrast, the fraction of neurons with MI on the second previous *d*_1_ (green) decreased rapidly throughout the current trial. There was also a fraction of neurons with significant MI about the second interval (*d*_2_) in the current (Fig. 4d, red lines), previous (blue) and second previous (green) trials.

These results differ from those obtained with recordings in PPC which showed that neurons only had information about the value of stimuli in previous trials [6]; information about the current one supposedly is processed in other areas. Our model assumes an area in which information about the current and preceding stimulus has converged.

### Ideal observer model

We verified above that the candidate causes for the contraction bias are indeed present in the neuronal activity of the trained networks. Now, taking advantage of the fact that normative models can help identify which computations are involved in producing the observed behavior, we formulate a model containing the possible causes of the bias. In doing so, we expect to understand how those factors operate and interact to generate the bias. To model behavior, we took the viewpoint of an ideal observer who makes noisy measurements of the intervals *d*_1_ and *d*_2_, has knowledge about the stimulus set, has access to the sensory history and can construct a belief about the duration of the presented interval. Measurements of the two time intervals are assumed to be Gaussian with means *d*_1_ and *d*_2_ and standard deviations that obey the scalar law (*σ*_1_ = *W*_1_*d*_1_ and *σ*_2_ = *W*_2_*d*_2_, respectively, where the parameters *W*_1_ and *W*_2_ are the corresponding Weber fractions. Methods) [44], [45]. We start by describing the sensory history of the first interval. The observation (*o*_1,*n*_) made at trial *n* combines the noisy measurement of the current interval 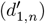 with the measurements of a few (*m*) recent past intervals, and with the observation made at a longer temporal scale (*o*_1,*n−m−*1_). That is,

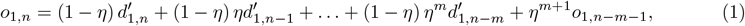

where *η* weights the contribution of preceding intervals (see Methods for further details) and 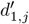(*j* = *n* − 1, …, *n* − *m*) is the noisy measurement of the first interval presented in trial *j*. We assumed that the long-term sensory history, *o*_*n−m−*1_, is in a Gaussian stationary regime, with its mean ℳ_*st*_ and variance 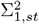 obtained self-consistently (Methods). With these assumptions, the observation of the interval *d*_1_ turns out to also be a Gaussian variable (𝒪_1_), with mean ℳ and variance 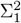. The observation of the second interval is simply the noisy measurement 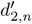 (for a model that considers the sensory history of the second interval see Supplementary Text S1). The prior probabilities of *d*_1_ and *d*_2_ were assumed to be discrete (Fig. 1b). Decisions, “*d*_1_ *> d*_2_” or “*d*_1_ *< d*_2_”, were made from the posterior probabilities using the *maximum a posteriori* decision rule (Methods). The general model in Eq. (1) depends on only three parameters: *W*_1_, *W*_2_ and *η*. We can now determine the observation probability of the first interval in the current trial *n*. Although this can be done for an arbitrarily long transient history (any *m*. Methods), in this work we will consider only the versions of the model with *m* = 0 and *m* = 1. The reason for not taking larger values of *m* is that the population with significant information about *d*_1,*n−*2_ is of a smaller size than the populations with significant information about the intervals in the current and the immediately preceding trials (Fig. 4c). For *m* = 0 the observation probability has mean and variance

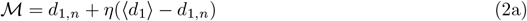

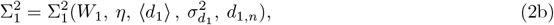

where ⟨*d*_1_⟩ is the first moment of *d*_1_ in the stimulus set and 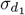 is its variance (see Methods for the full expression of 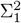). Note that both ℳ and Σ_1_ could contribute to the contraction bias: The mean indicates explicitly that the first interval is contracted toward the interval mean, by an amount proportional to the difference between the true value of the interval and the *d*_1_ mean. The variance provides a Bayesian explanation of the bias, which operates when the uncertainty in the first interval exceeds the uncertainty in the second interval, i.e., when Σ_1_ *> σ*_2_ [15]. Interestingly, the sensory history parameter *η* modulates the variance contribution to the bias (Methods). For each network, the parameters *η, W*_1_ and *W*_2_ were fitted to optimize the similarity between the accuracy curves of the model and the network (Methods). The fit for the example network 1 is shown in Fig. 5a (see Extended Data Fig. 3a for example network 2) and the network root-mean-square error (RMSE) distribution of the fits appears in Fig. 5b. A measure of the contraction bias, *B*_*fit*_, obtained from the fitted performance, is given in Fig. 5c.

**Fig. 5.**
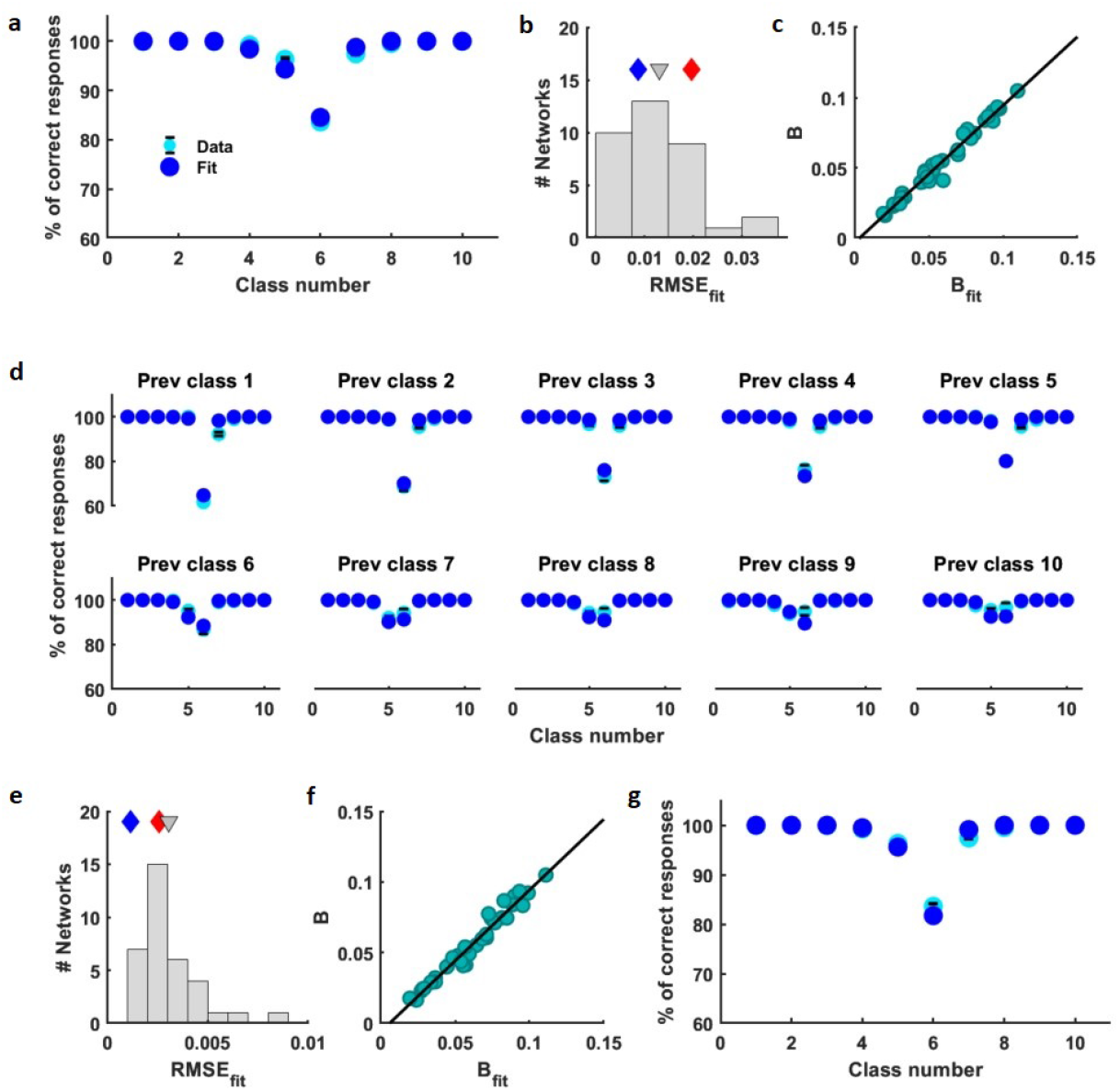
Bayesian model fits. **a-c**. Fit with model version *m* = 0 (Eqs. (2a,b)). **a**. Fit of percentage of correct test trials (computed on the diagonal block) of example network 1: behavioral data (cyan dots with error bars are the data in Fig. 1e, right) and fit (dark blue circles). **b**. Histogram of RMSEs (Root Mean Square Error from the difference between the behavioral data performance and the model-fitted performance) over 35 networks. Histogram’s mean is represented by gray triangle. Diamonds indicate the RMSEs values of network 1 (blue) and network 2 (red). **c**. Correlation between the phenomenological bias (*B*) and the bias from the model fit (*B*_*fit*_), computed over 35 networks (correlation=0.98; p-value=5.6e-27). Both *B* and *B*_*fit*_ were computed with Eq. (14) (Methods). **d-g**. Fit with model version *m* = 1 (Eqs. (3a,b)). **d**. Fit of the ten accuracy curves conditioned over each possible preceding diagonal class of example network 1 (cyan circles are the behavioral data in Fig. 2a). **e**. Histogram of RMSEs of performance fits over 35 networks. **f**. Correlation between *B* and *B*_*fit*_ (correlation=0.97; p-value=4.2e-26). **g**. Marginal accuracy curve for network 1, obtained by marginalizing over the preceding first interval both the data and fit in panel (d) (see Methods).

The model version *m* = 0 considers the effect of the long-term sensory history on the observation of the first interval, but neglects the contribution from recent trials. To examine this issue, we now turn to the case *m* = 1; the mean and variance of the observation probability are

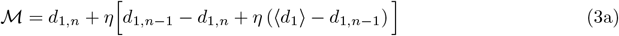

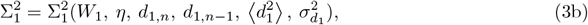

where 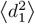 is the second moment of *d*_1_ and 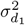 is its variance (Methods). The observation of the current interval contracts towards *d*_1,*n−*1_ which, in turn, appears contracted towards ⟨*d*_1_⟩. Again, there is a Bayesian source for the bias when Σ_1_ *> σ*_2_. Note that the preceding interval affects the observation variance and, hence, short-term sensory history modulates the Bayesian contribution to the bias. Thus, in the model, all the sources of the contraction bias appear to be intricately related.

To fit the model parameters (*η, W*_1_ and *W*_2_), we considered the 10 accuracy curves that result from conditioning on each of the possible preceding classes (Fig. 2a for the example network 1 and Extended Data Fig. 1c for the network 2), and used them all to define the similarity between model and data (Methods). The fits for the example networks 1 and 2 are shown in Fig. 5d and Extended Data Fig. 3b, respectively. The network distribution of the RMSEs of the fits is in Fig. 5e, and the relationship between the phenomenological bias *B* and the bias quantified from these fits appears in Fig. 5f. The average accuracy for each current class, obtained by marginalizing over the preceding first interval (Methods and Supplementary Text S2), is in Fig. 5g (Extended Data Fig. 3c for network 2).

The model in Eq. (1) describes the sensory history of the first interval. However, the activity during the delay period also has information about the second interval in the preceding trial, *d*_2,*n−*1_ (Fig. 4d). A more general model that considers the combined sensory history of *d*_1_ and *d*_2_ is provided in Supplementary Text S1. Surprisingly, the quality of the fit with this model is identical to that obtained with the model in which the contribution of the second interval is not considered (Extended Data Fig. 4 and Supplementary Table S1). Since the inclusion of the second interval did not improve the fit, for the sake of simplicity, in the following we will not take it into account.

Before we showed that a weak contraction bias is still present in networks trained resetting the network activity at the end of each trial (Fig. 2c). We now wonder about the model that describes this situation, and about the origin of the bias in these networks. Since the current trial is not affected by the sensory history, the observation model in Eq. (1) should be excluded. Instead, we considered the possibility that neural circuits may have knowledge of the stimulus statistics (e.g., knowledge of ⟨*d*_1_⟩), which presumably could have been learned during training and kept stored in the network synapses. We then contrasted the hypothesis that the circuits have knowledge about ⟨*d*_1_⟩ with the hypothesis that they do not. In the first case the observations have mean ℳ = *d*_1,*n*_ + *μ*(⟨*d*_1_⟩ − *d*_1,*n*_) and variance 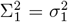, while in the second these are ℳ = *d*_1,*n*_ and 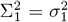, respectively. The parameters of the first model are *μ* and *σ*_1_ (but note that *μ* is not a sensory history parameter), while the only parameter of the second model is *σ*_1_. We fitted the behavioral data for each network with both models obtaining comparable results in terms of the fit quality (Extended Data Fig. 5). Comparison of the two models using BIC or AIC tends to favor the model that does not require knowledge of ⟨*d*_1_⟩ (Supplementary Table S2). This means that the Bayesian explanation could be enough to account for the contraction bias in networks trained with activity reset, without the need to store the interval mean in the synapses [29].

### Correlations between the phenomenological bias and the model parameters

We now use the normative model to examine how the sensory history and the uncertainty in the observations operate and interact to generate the bias. Since the variance of the observation of *d*_1_ depends on sensory history, this analysis is not immediate. First, we noted that the condition Σ_1_ *> σ*_2_ determines whether the uncertainty in the observations contributes to the bias [15]. We will say that in networks where this condition is satisfied the Bayesian contribution to the bias is operative. Then, we sorted the 35 networks into two groups: a group in which that condition was satisfied (the positive group) and a group in which it was not (the negative group). To obtain further insight into the role of each parameter, for each network group we evaluated the correlation between the phenomenological measure of the bias (*B*) and the model parameters: the sensory history parameter (*η*) and the two observation uncertainties (Σ_1_ and *W*_2_). Note the use of Σ_1_ instead of *W*_1_, this is because the quantity relevant for determining the bias is the total variance of the observation probability of *d*_1_.

We start with the simplest version of the model (*m* = 0, Eqs. (2a) and (2b)). About half of the networks (18/35) belonged to the positive group (Σ_1_ *> σ*_2_), signaling a contribution from observation uncertainty to the contraction bias. As expected, in positive networks *B* was significantly and positively correlated with Σ_1_ (Fig. 6a, left and Supplementary Fig. S4a). Instead, *B* was not significantly correlated with *η*(Fig. 6a, middle). This is surprising, given that the mean ℳ explicitly describes the mechanism by which the sensory history contributes to the formation of the bias, Eqs. (3a) and (3b). The explanation for this paradox is that the sensory history also contributes to Σ_1_, and does so in such a way that *η* is negatively correlated with the variance (Fig. 6b and Supplementary Fig. S5). This reduces the overall effect of *η* on the bias. Finally, *B* was not significantly correlated with *W*_2_ (Fig. 6a, right), a parameter that only becomes relevant after the end of the delay period. We observed that in most networks the difference between the noise parameters, *W*_1_−*W*_2_, was positive (Fig. 6c). In negative networks the situation was the other way around: *B* and Σ_1_ were not significantly correlated (Fig. 6d, left, and Supplementary Fig. S4b), in contrast, *B* and *η* were positively correlated (Fig. 6d, middle). In negative networks *B* and *W*_2_ were also not significantly correlated (Fig. 6d, right).

**Fig. 6.**
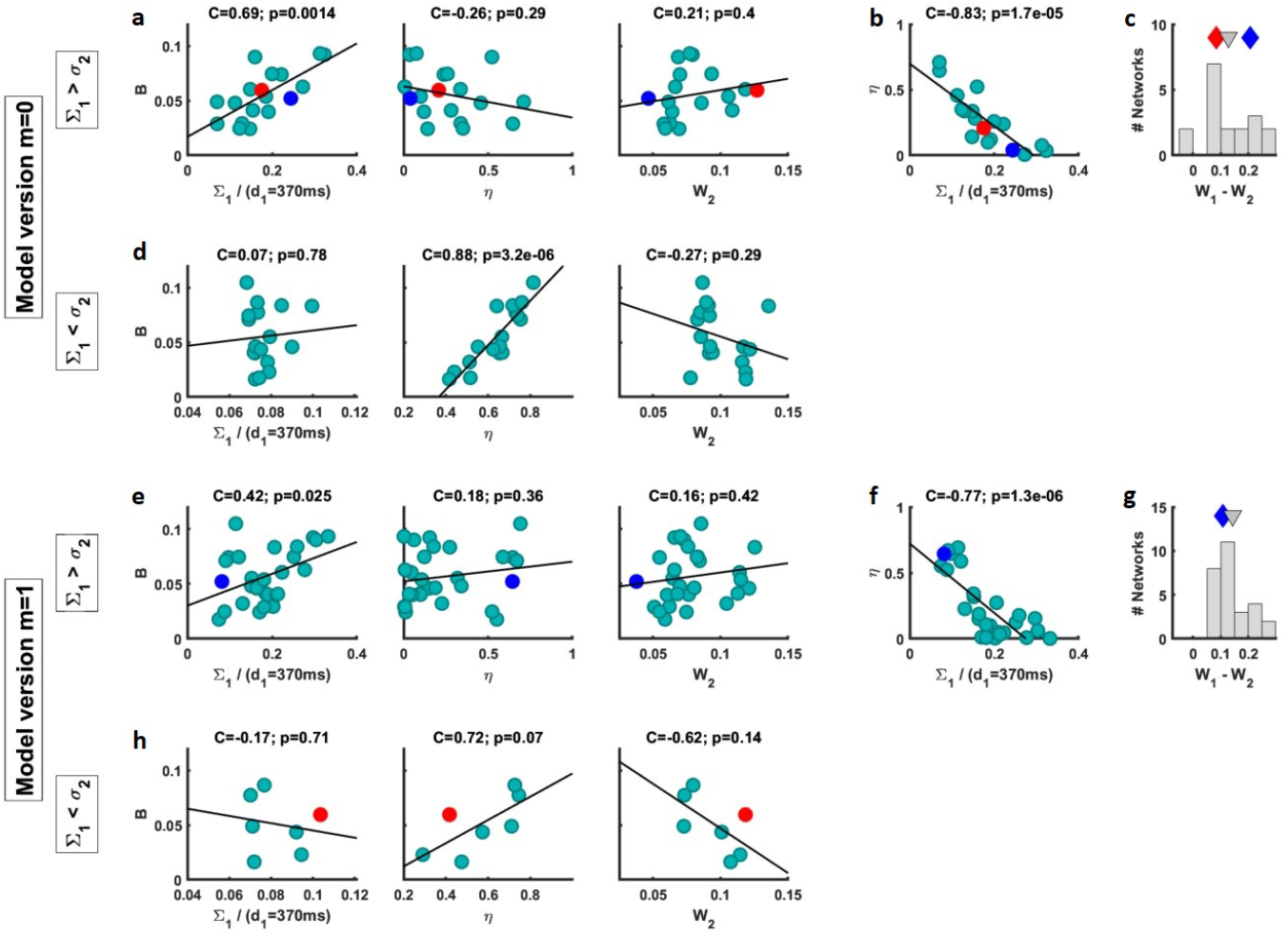
Correlations of parameters from both model versions *m* = 0 and *m* = 1, sorted by network type (Σ_1_ *> σ*_2_ or Σ_1_ *< σ*_2_). **a-c**. Correlations over the 18 networks classified as positive (Σ_1_ *> σ*_2_) by model version *m* = 0. **a**. Correlations between the bias *B* and the model parameters. **Left**: Correlation between *B* and the total variance of the observation probability (Σ_1_) for *d*_1_ = 370 ms (for other values of *d*_1_ see Supplementary Fig. S4a). **Middle**: Correlation between *B* and the sensory history parameter (*η*). **Right**: Correlation between *B* and the noise parameter of *d*_2_ measurements (*W*_2_). **b**. Correlation between *η* and Σ_1_ for *d*_1_ = 370 ms (see Supplementary Fig. S5). **c**. Histogram of the difference (*W*_1_ − *W*_2_). Gray triangle indicates the histogram’s mean. **d**. Correlations between *B* and the model parameters from 17 networks classified as negative (Σ_1_ *< σ*_2_) by model version *m* = 0. **Left**: Correlation between *B* and Σ_1_ for *d*_1_ = 370 ms (see Supplementary Fig. S4b). **Middle**: Correlation between *B* and *η*. **Right**: Correlation between *B* and *W*_2_. **e-g**. Correlations over the 28 networks classified as positive by model version *m* = 1. **e**. Correlations between *B* and the model parameters. **Left**: Correlations between *B* and Σ_1_ for the current and previous first stimulus equal to 370 ms (see Supplementary Fig. S6). **Middle**: Correlation between *B* and *η*. **Right**: Correlation between *B* and *W*_2_. **f**. Correlation between *η* and Σ_1_ for the current and previous first stimulus equal to 370 ms (the rest of the cases hardly change). **g**. Histogram of *W*_1_ − *W*_2_. **h**. Correlations between *B* and the model parameters from 7 networks classified as negative by model version *m* = 1. **Left**: Correlation between *B* and Σ_1_ for the current and the previous first interval equal to 370 ms (see Supplementary Fig. S7). **Middle**: Correlation between *B* and *η*. **Right**: Correlation between *B* and *W*_2_. Blue markers refer to network 1 and red markers refer to network 2. C indicates correlation values and p is the p-value.

A similar pattern of correlations held for the model version with *m* = 1, Eqs. (3a) and (3b). For most networks (28/35), the fit satisfied that Σ_1_ *> σ*_2_, meaning that the Bayesian contribution to the bias was present more often than in the simpler version. Again, in positive networks the bias *B* was favorably correlated with Σ_1_ (Fig. 6e, left and Supplementary Fig. S6). Instead, its correlation with *η* was not significant (Fig. 6e, middle). The reason for this feature was again the competition between the contributions of the sensory history to the mean ℳ and to the variance 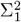 (Fig. 6f). Again, *B* was not significantly correlated with *W*_2_ (Fig. 6e, right). Finally, the difference *W*_1_ − *W*_2_ was still positive (Fig. 6g). As for the negative networks, the model version *m* = 1 gives predictions similar to those found with the version *m* = 0 (Fig. 6h and Supplementary Fig. S7), although the positive correlation between *B* and *η* was somewhat below significance level (Fig. 6h, middle).

In summary, inspired by data taken from the trained sRNNs, we began by proposing a normative model, Eq. (1), which included the factors supposedly responsible for the contraction bias, namely, the sensory history and the observation uncertainty relative to *a priori* knowledge. Fitting the data with this model confirmed that both factors were necessary: The measurement of *d*_1_ is affected by a fixed noise (*W*_1_), yet, the uncertainty of its observation is also influenced by the sensory history. We then conclude that the contraction bias results from the sensory history, the uncertainty in interval measurements, and an interaction between these two factors.

### State-space geometry is related to behavior

So far we have analyzed behavior and neural activity separately. Guided by a normative model of behavior we have 1) distinguished two groups of networks in which the Bayesian contribution to the bias would have different relevance and 2) noted that the sensory history affects the Bayesian contribution to the bias. Nevertheless, those analyses do not inform us about the neural mechanisms that could implement the computations suggested by the model. To examine this issue, we now investigate how state-space geometry relates to behavior. The task requires that during the presentation of *d*_1_ the neural activity represents an estimate of its duration and then maintains it during the delay period, until the end of this interval. Accordingly, we expect that over these two epochs an estimate of *d*_1_ may be represented in the geometric properties of the trajectories [20], [21]. We have seen that during interval presentation the trajectories define a one-dimensional manifold characterized by its curvature (Fig. 3a) while after the completion of each interval (SF1) the trajectories remain distant, reaching a quasi-attractor region at the end of the delay epoch (Figs. 3b,c). So, we will now assume that estimations of *d*_1_ are represented, first, in the curvature of the manifold (the “curved manifold hypothesis” [21]) then, during the delay period, in the relative distances of the trajectories and, finally, in the distances between the terminal states on the quasi-attractor. However, the nature of the estimator is unknown, since in delayed comparison tasks the estimate of the first stimulus is not reported, only the comparison choices are available. This contrasts with magnitude reproduction tasks, in which the reproduced magnitude can be taken as a noisy version of the estimate [11]. Given the limited information about what estimator of *d*_1_ employed the networks, we considered, apart from the true *d*_1_, two other alternatives, the Bayesian and Maximum Likelihood (ML) estimators (*d*_1,*Bayes*_ and *d*_1,*ML*_, respectively. Methods), and studied how they related with curvature, relative distances between neural trajectories and distances between terminal states on the attractor.

To find out which estimator best explained the curvature in the neural activity, we projected the neural states in the curved manifold onto an encoding vector **u**, defined as the unit vector pointing from the state associated with the shortest to the longest *d*_1_ in the set [21] (Methods). For the example network 1 (Fig. 7a) the distances *P*_*j*_ between the projections (See inset; *j* = 1, …, 5 labels the values of *d*_1_ in the diagonal block) gave a warped representation of *d*_1_. We then asked whether the *P*_*j*_ more closely reflected the true *d*_1_ values or one of the two estimates, *d*_1,*Bayes*_ or *d*_1,*ML*_. To confront these possibilities, we first separately fitted a linear expression (*aP*_*j*_ + *b*) to the true *d*_1_ and to the two estimators (the three regressions appear in Fig. 7b). Then, we compared the corresponding goodness of fit using their RMSEs (Fig. 7c); for this network the test favored the Bayesian estimator hypothesis, and the same happened for most positive networks (Extended Data Figure 6a). To determine which estimator of *d*_1_ best accounted for the relative distances between the neural trajectories during the delay epoch, we followed a similar approach. The relative distance *D*_*j*_(t) between each trajectory *j* = 1, …, 5 and the reference trajectory *d*_1_=370 ms, was defined following the KiNeT method [19] (Methods). The time courses of these distances for the example network 1 (Fig. 7d) and their temporal means *D*_*j*_ ≡ ⟨*D*_*j*_(*t*)⟩ (inset), also appeared warped as a function of *d*_1_. We then asked whether *D*_*j*_ coded better the true *d*_1_ values or their Bayesian or ML estimators. To confront those alternatives, we separately fitted a linear expression (*aD*_*j*_ + *b*) to these three estimates (Fig. 7e) and we compared their RMSEs [20] (Fig. 7f). Again, for this network the test favored the Bayesian estimator hypothesis (see also Extended Data Fig. 6b). Finally, to test the estimators on the quasi-attractor we computed the distances *δ*_*j*_ between terminal points of the trajectories on the attractor region and made an analysis similar to the two previous ones. Again, the test favored the Bayesian estimator (Figs. 7g-i and Extended Data Fig. 6c). The results of the three tests provide us with links between behavior and state-space geometry, for those networks in which the bias can be explained by Bayesian arguments.

**Fig. 7.**
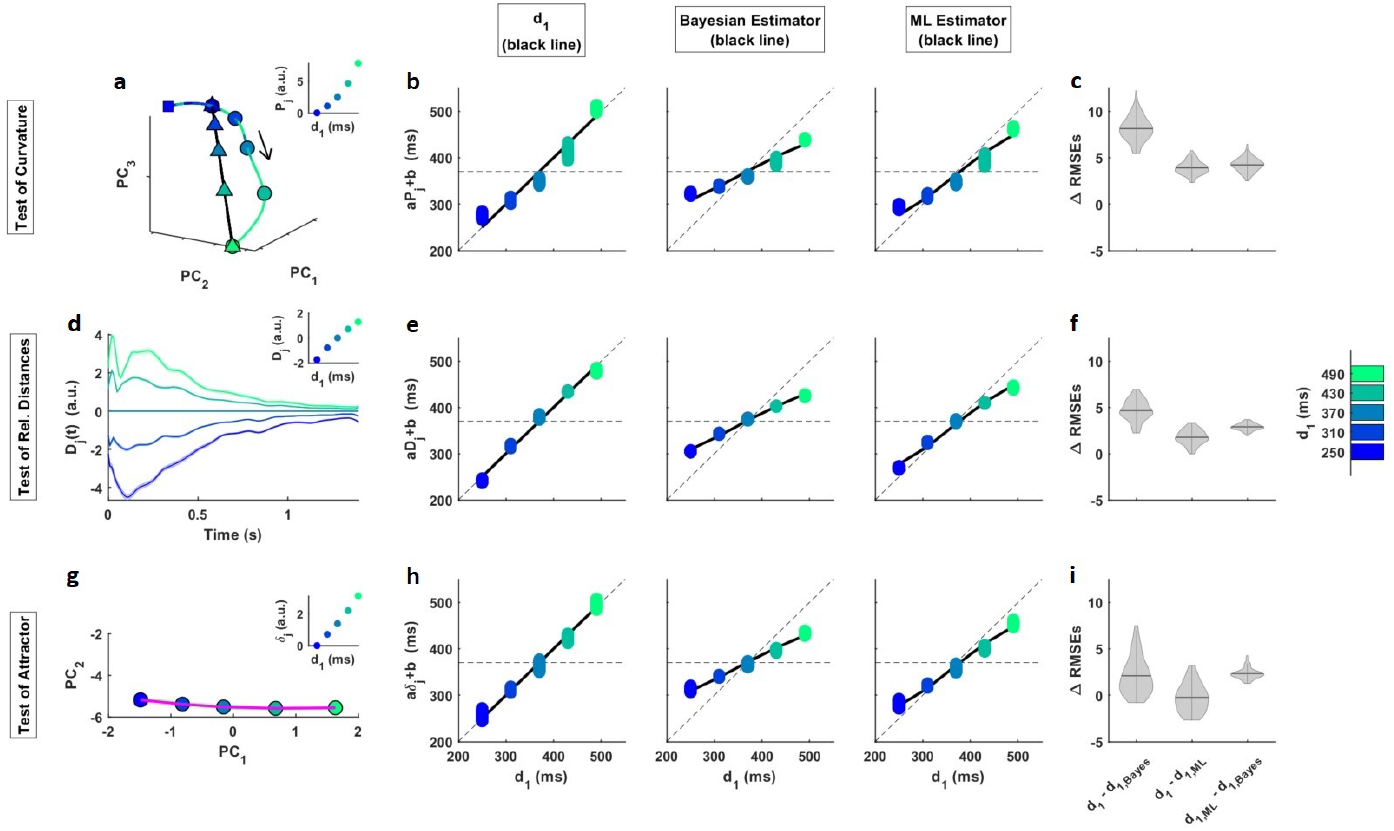
Several geometric features code the Bayesian estimator of *d*_1_. **a-i** Example network 1. **a-c**. Curvature of the one-dimensional curved manifold during the presentation of the first stimulus. **a**. Neural trajectories and projections (triangles) of each SF1 terminal state (circles) onto the encoding axis, **u** (black line). Squares represent the beginning of the figure, located within the first stimulus. Neural trajectories are embedded within a state space built with the first 4 PCs, which explain 97% of variance. **Inset** shows, as a function of *d*_1_, the Euclidean distances *P*_*j*_ (*j* = 1, …, 5 labels the values of *d*_1_ in the diagonal block) between projections, taken as reference the projection of the shortest *d*_1_. **b**. Linear regression of the distances between projections (*aP*_*j*_ + *b*) to the true *d*_1_ (left), the Bayesian estimator (middle) and the ML estimator (right), as a function of *d*_1_. Colored dots show multiple estimates of the regression derived from bootstrapping (*n*=100). Solid lines indicate the true *d*_1_ (left), the Bayesian estimator (middle) and the ML estimator (right) versus *d*_1_. **c**. Violin plots (kernel probability density) on shufflings of the RMSE differences between the three hypotheses (*d*_1_, Bayesian estimator and ML estimator). The width of the gray area indicates the density of the shufflings located there. Horizontal line inside of each violin plot indicates the distribution’s mean. The chosen hypothesis is always the Bayesian estimator **d-f**. Relative distances during the delay period. **d**. Relative distance, *D*_*j*_(*t*), between each trajectory *j* = 1, …, 5 and the reference trajectory *d*_1_=370 ms. Neural trajectories are embedded in a state space built with 6 PCs, which explain 92% of variance. Shadings indicate the mean ± 95% CIs computed from 100 bootstrap resamples. **Inset** shows the temporal means of relative distances as a function of *d*_1_. **e**. Linear regression of the mean relative distances (*aD*_*j*_ + *b*) to the true *d*_1_ (left), the Bayesian estimator (middle) and the ML estimator (right) as a function of *d*_1_ (same format as in (b)). **f**. Analogous to (c), but for the relative distance analysis. The preferred hypothesis is always the Bayesian estimator. **g-i**. Distances between terminal states on the quasi-attractor. **g**. One-dimensional quasi-attractor region (magenta line), at the end of the delay period, embedded within the first 2 PCs space (99.7% of variance). **Inset** shows the Euclidean distances, *δ*_*j*_, between two *d*_1_ terminal states (taken the shortest *d*_1_ as reference) on the attractor region, as function of *d*_1_. **h**. Linear regression of the terminal states on attractor (*aδ*_*j*_ + *b*) to the true *d*_1_ (left), the Bayesian estimator (middle) and the ML estimator (right) as a function of *d*_1_ (same format as in (b)). **i**. Analogous to (c), but for the terminal states analysis. The attractor was analyzed with a time window of size 100 ms at the end of the delay period. In the vast majority of cases, the Bayesian estimator is chosen.

Earlier we noted that, according to Bayesian arguments, the contraction bias increases when noise is introduced into the trained network (Fig. 2b, right). Based on the relationship between behavior and geometry just described, we predict that increasing noise levels would lead to greater distortion of the affected geometric features. Our analysis confirmed this expectation. Specifically, when noise was applied during stimulus presentation, the distances *P*_*j*_ exhibited a more warped representation compared to the noiseless case (Extended Data Fig. 8a). We observed a similar effect on the relative distances *D*_*j*_ when noise was introduced during the delay period (Extended Data Fig. 8b). The results in Figures 7 and 8 reinforce the idea that Bayesian computations already take place at the sensory stage of the task and extends to the time of stimulus comparison.

Given that network 2 does not satisfy Σ_1_ *> σ*_2_ we do not expect it to pass the tests well. To confirm this expectation, we conducted similar analyses on network 2. Despite the presence of curvature in the 1-dimensional manifold, the Bayesian estimator did not outperform the other estimators in explaining the neural trajectory geometry in this network (Extended Data Fig. 7).

Previously, we quantified the contraction bias in terms of the discrimination curves (*B*). Now, we can quantify it using the estimator *d*_1,*Bayes*_, derived from the behavioral fit (Methods). This gave us a measure for the bias (*B*_*Bayes*_) that correlated well with the empirical measure *B*(C=0.96, p*<*10-6). In addition, *B*_*Bayes*_ had the same pattern of correlations with the model parameters as *B*(Supplementary Fig. S8).

### Effect of sensory history on state-space geometry

We saw that the sensory history modulates the contraction bias (Fig. 2a). So, according to the results in Fig. 7, it can be expected to modulate geometry as well, and to do so in a way consistent with the change it produces on the bias. For example, in network 1, when varying the previous class along the set’s upper diagonal (i.e., from class 1 to 5. Fig. 1b), the bias decreases approximately by half (*B*=0.12 vs. 0.06. Fig. 2a). Therefore, if our expectation were correct, the geometric features should exhibit a more warped representation of the current *d*_1_ in trials preceded by class 1 than in trials preceded by class 5. To verify this, we first checked whether the geometry of the trajectories still codes the Bayesian estimator when conditioning on the class of the preceding trial. The tests for the curvature, relative distances and positions on the quasi-attractor, obtained conditioning separately on the previous classes 1 or 5, showed that *d*_1,*Bayes*_ was still the preferred estimator (Extended Data Fig. 9). Next, we related each of the three geometric features with the contraction bias, defining the compressions *C*[*Z*] (*Z* = *P*_*j*_, *D*_*j*_ or *δ*_*j*_) of the associated *d*_1_ representations (Methods, Eq. 52). Finally, we computed each *C*[*Z*] in the two conditions (previous class 1 or 5). In all three cases the compression was higher when the preceding class was class 1. The compressions from the curved manifold were *C*[*P*] = 0.49 (previous class 1) and *C*[*P*] = 0.42 (previous class 5) with p-value=0.004 (Mann-Wilcoxon test over two bootstrap distributions, each with 100 resamples). For the mean relative distances, *C*[*D*] = 0.33 (class 1) and *C*[*D*] = 0.30 (class 5), p-value=0.09. Finally for the positions on the attractor, *C*[*δ*] = 0.43 (class 1) and *C*[*δ*] = 0.21 (class 5), p-value=6e-21. The decrease in significance during the delay period and its important recovery at the end of this epoch could be related to the initial decrease and subsequent rise in the information about the previous intervals that occur during the memory period (Fig. 4c,d). Network 2 has a similar bias *B* as network 1 and undergoes a similar change in bias when conditioning on prior classes 1 or 5 (*B*=0.14 vs. 0.06. Extended Data Fig. 1c). However, since this network does not satisfy that Σ_1_ *> σ*_2_, *d*_1,*Bayes*_ was not unambiguously the preferred estimator (data not shown).

## Discussion

The contraction bias is influenced by the stimulation history [6], [26]. However, it is not solely explained by the action of preceding stimuli; it can also arise from integrating prior knowledge with noisy observations of the stimuli [9], [15], [46]. In this study, we aimed to unravel how the contraction bias emerges from the interplay of these factors by training sRNNs on a delayed comparison task. Through the analysis of behavioral and firing activity data, we successfully identified the sources of the bias and proposed a normative model that elucidated the network computations responsible for this phenomenon. Interestingly, the majority of trained networks (80%) exhibited a Bayesian explanation for the contraction bias, where it played a prominent role while sensory history modulated the uncertainty in the observation of the first stimulus. Although sensory history was consistently present, it was only correlated with the bias in a minority of networks (20%).

During the task execution, we observed successive geometric properties in the state-space trajectories, which exhibited compressed representations of the duration of the first interval. Specifically, this compression occurred during the presentation of the stimulus, where it was reflected in the curvature of a 1-dimensional manifold. Subsequently, during the delay period, the compression was observed in the relative distances between the trajectories. Finally, towards the end of that epoch, it manifested in a one-dimensional quasi-attractor. Importantly, these compressions aligned with the predictions of the Bayesian estimator derived from our normative model, indicating that the contraction bias arose from the geometric characteristics of the trajectories in state space.

Several aspects of our results can be compared with recent experimental studies of neural activity in both delayed comparson tasks and reproduction tasks. Akrami et al. found that during a discrimination task neurons in PPC carried information about recent sensory history [6]. However, this area did not present information about the stimulus of the current trial, so it does not correspond to the model considered in our work, which assumes an area that has information about current and preceding stimuli. On the other hand, in a temporal reproduction task Sohn et al. found that a curved one-dimensional manifold exhibited a compressed representation of the time interval, that was matched by the compression predicted by the Bayesian estimator [21] (see also [47]). Our results showed that a similar relationship between population activity and behavior may occur even when an estimate of the interval is not reported in the task.

In psychophysics experiments [15], [48], fMRI studies [28] and electrophysiology recordings [25], aimed at investigating contraction bias, the question has arisen as to at what trial epoch and in which brain areas the prior is combined with noisy sensory information. In the study by Ashourian et al. (2011), it was hypothesized that the Bayesian operation is performed explicity, finding that it occurs during the comparison period or later. Our results challenge this hypothesis by demonstrating that the Bayesian estimator is consistently manifested in the geometry of the state space throughout the trial, at least until the decisional stage. Furthermore, we observed that the a priori information was not explicitly combined with sensory information; instead, the dynamics of network activity directly transformed the duration of the first interval into its Bayesian estimator. We agree with [28] in that prior information about the set is already used during the stimulation epoch. Benozzo and Genovesio (2013) focused their study on the decisional stage while our research emphasized the stimulation and maintenance epochs of the task. Exploring the dynamics of stimulus comparison and beyond can contribute valuable insights to the field, potentially necessitating novel approaches and methodologies. The analysis of the time interval discrimination task presented here can be adapted for use in monkeys or rodents, and we anticipate that the results will also apply to other delayed comparison tasks, including discrimination of auditory [6], [49] or tactile stimuli [50], [51]. Our work may inspire the adaptation of such experiments to verify our findings. In addition, our use of spiking RNNs offers practical advantages, such as the possibility of a straightforward application of the analysis techniques developed here to real recordings of neural activity.

Our trained spiking RNNs exhibited human and animal-like behavior. For example, in agreement with psychophysics experiments conducted in humans [15], the networks showed increased bias when perturbed with noise. In contrast, other training procedures yielded network models that failed to reproduce the biases exhibited by behaving animals [31], [52]. The question of how neural networks should be trained to obtain models consistent with observed behavior has received some attention [53], but is not yet well understood. One possible approach is the use of noise; indeed, it helped to obtain animal-like behavior [30] and to account for biases due to temporal expectation [4] with trained RNNs of rate neurons. However, networks of noisy GRUs optimized in a frequency discrimination task achieved perfect performance and thus did not exhibit the contraction bias [31]. A possible cause of such good performance of the networks trained in [31], [52] is the use of neuron and network models with computational capabilities that are beyond those of real brains, and thus can outperform animal behavior, even in the presence of noise [31]. Although our training technique is not biologically realistic, the resulting spiking RNNs are plausible. Given the usefulness that RNNs are showing in understanding how cognitive tasks can be solved in biological networks, fully understanding how factors such as the neural model or pre-training procedures lead to models in agreement with animal behavior is a crucial problem that deserve further research.

This work advances the understanding of the origin of the contraction bias by proposing a unified description of the potential sources of this bias and showing that the Bayesian explanation is prevalent in recurrent networks trained to discriminate temporal intervals. The results raise the question of whether they occur in the brain; in this regard, one of the most pressing questions is the elucidation of in which areas and in what way the contraction bias is generated. Existing recordings obtained from monkeys during the tactile discrimination task, initially conducted to investigate the evolution of perceptual decisions in sensory and frontal areas [51], [54], offer a valuable opportunity to undertake the mentioned study.

Our work emphasizes the significance of employing trained networks to establish connections between behavior and neural activity, particularly in unraveling how perception is influenced by sensory history, short-term memory and long-term plasticity. By doing so, our methods can help to bridge the gap between psychophysics [12], [15], [48], [55] and electrophysiology [6], [23], [25], or other techniques used to measure brain activity [28]. Furthermore, our approach extends beyond delayed comparison tasks such as time interval estimation [11], [18], magnitude estimation [12], [13], or delayed response tasks [24], [29], [56], [57]. In these instances, we anticipate that our methods will aid in determining in which way and in which trial stage sensory information and prior knowledge combine to give rise to perception. In summary, our research improves our understanding of perception by uncovering the interplay between current and recent stimulation, prior knowledge, and contextual cues, which lead to perceptual biases. We emphasize the central role of Bayesian principles and challenge existing assumptions about how the integration with prior information occurs. Furthermore, we elucidate the influence of recent sensory history on perceptual bias as a modulation of Bayesian computations.

## Methods

### Temporal Interval Discrimination (TID) task

Spiking Recurrent Neural Networks (sRNNs) were trained to decide which of two time intervals (*d*_1_, *d*_2_), selected from the stimuli set (Fig. 1b) and presented sequentially separated by a delay epoch, was the shortest. The task scheme (Fig. 1a) began with the Start Cue event. After a 1s-delay, the first time interval (*d*_1_) was presented (SO1 and SF1 are the stimulus onset and offset of this interval, respectively). A second time interval (*d*_2_) of variable duration (Fig. 1b) was displayed after a 1.5 s delay period (SO2). Both *d*_1_ and *d*_2_ were introduced as step signals. In training trials, one second after the offset of *d*_2_ (SF2) a target signal, *z*_*tgt*_(*t*), was presented for a time *T*_*tgt*_ (0.5 s). This target was a beta distribution whose sign indicated the correct decision, i.e., positive for decision *d*_1_ Longer, *L* ≡ “*d*_1_ *> d*_2_”, and negative for decision *d*_1_ Shorter, *S* ≡ “*d*_1_ *< d*_2_”. Finally, the duration of the inter-trial-interval (ITI) was variable and it was defined so that all trials have the same duration (5.5 s).

### Network model

We considered sRNNs of *N*=500 leaky integrate-and-fire (LIF) neurons. We will use bold characters to denote arrays and subscripts to denote array elements. The dynamics of the membrane potential of a spiking neuron *i*(*i* = 1, …, *N*), *V*_*i*_(*t*), is

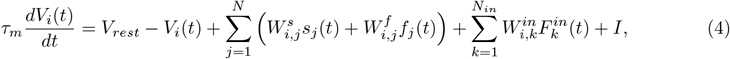

where *τ*_*m*_ = 10 ms is the membrane time constant and *V*_*rest*_ = −65 mV is the rest potential. Once the membrane potential reaches a threshold value (*V*_*th*_ = −55 mV) an action potential (spike) is produced. Immediately after an action potential, the membrane potential is reset to *V*_*reset*_ = *V*_*rest*_. Whenever a neuron *j* fires a spike, its corresponding synaptic currents, *s*_*j*_(*t*) and *f*_*j*_(*t*), are incremented by 1, otherwise

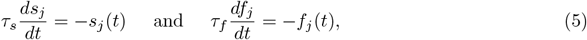

where *τ*_*s*_ = 100 ms for the slow current, *s*_*j*_(*t*), and *τ*_*s*_ = 5 ms for the fast current, *f*_*j*_(*t*). Slow (fast) current flows from a pre-synaptic neuron *j* to a post-synaptic neuron *i* through the hidden recurrent connections 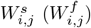. An input layer of *N*_*in*_ = 2 neurons introduces both stimuli (*d*_1_, *d*_2_) into the network as external input signals, 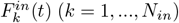, through the input weights 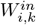, defined from a uniform distribution. The current *I* consists of two terms, a contribution evaluated before learning by applying the recurrent targets as external inputs and a global inhibition of intensity 0.3.

### Full-FORCE training algorithm

The full-FORCE algorithm was used as a target-based supervised learning method to train sRNNs [36]. The full-FORCE scheme trains a sRNN (called task-performing network) to generate the desired target output, *z*_*tgt*_(*t*), using target functions for the recurrent currents, 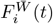 (where *i* = 1, …, *N*), obtained from an auxiliary RNN of rate neurons (known as target-generating network) (Supplementary Fig. S1). These target functions provide the necessary information about the currents to train the recurrent connections of the sRNN.

#### Target-generating network

The objective of this network of 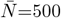 recurrently connected firing-rate units is to provide targets to the sRNN. Therefore, the target-generating network must be driven by the desired target signals, *z*_*tgt*_(*t*), in addition to the input events, ***F*** ^***in***^(*t*). The dynamics of the activity 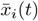 of neuron 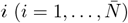 from the target-generating network is governed by

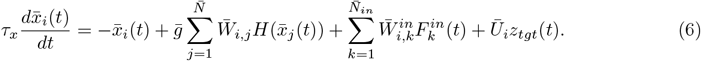

The non-plastic weights 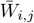 are built from a Gaussian distribution of zero mean and variance 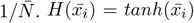 is the rate activity. 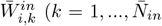 where 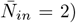 and 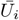 (only one output neuron is considered) are non-plastic synapses that introduce the input signal, ***F*** ^***in***^(*t*), and the target signal, *z*_*tgt*_(*t*), respectively, into the network. Both synapses were built from uniform distributions. Finally, 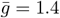 is a gain factor.

The auxiliary target functions, 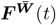, are transferred as targets to the task-performing network

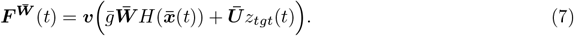

The 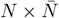 matrix ***v*** is used to solve a possible mismatch between the dimensions of 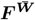, which is *N*, and of the target-generating network, which is 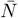.

*Training of task-performing networks and learning rule*. All the output weights ***W*** ^***out***^ and the post-synaptic weights (slow and fast) of *N*= 200 randomly chosen neurons were trained (Supplementary Fig. S1, red lines). The slow and fast currents flowing through the plastic recurrent synapses were concatenated in a 2*n*-dimensional vector ***c***(*t*) while the corresponding matrix ***W*** was an *N* x2*n* array. In each training trial both ***W*** and ***W*** ^***out***^ were adjusted to generate the optimal network response. Considering the auxiliary functions, 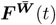, and the internal plastic currents, ***c***(*t*), the recurrent hidden weights were modified so that the following condition was met

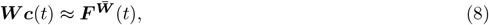

while for the output weights it was imposed that

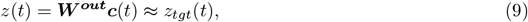

where *z*(*t*) is the output signal of the task-performing network given by an output layer of *N*_*out*_ = 1 neuron. The recurrent hidden weights were trained minimizing the cost function

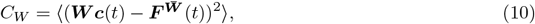

while the output synapses, ***W*** ^***out***^, were modified minimizing

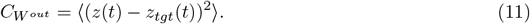

The optimization and update were done using the Recursive Least Squares Algorithm (RLS) [58]. *Test trials*. In test trials, a network output was considered correct if *z*_*tgt*_(*t*) and *z*(*t*) fulfilled the following condition during the presentation of the target [59]

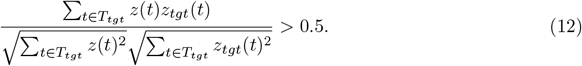

Conversely, a trial was taken as wrong if *z*(*t*) satisfied Eq. (12) with *z*_*tgt*_(*t*) replaced by −*z*_*tgt*_(*t*). In case *z*(*t*) did not satisfy any of these two conditions, then it was considered as a noisy response. These trials were a small fraction of the test trials and were not used in the analyses.

### Numerical experiments and simulation details

#### Training and simulations in the main numerical experiment

Thirty five different networks (obtained from different random initializations of the network parameters) were trained in a block of 5,000 training trials, presented in a continuous manner, without perturbing the network activity at the end of the trial. Each training trial followed the scheme in Fig. 1a. During the training stage, classes were selected randomly from Fig. 1b, excluding the central class (370, 370) ms. Once a network was trained, test trials were collected in the same way as the training trials, except that the target was not presented. During the testing stage, we considered three different trial blocks: the full block, the horizontal block and the diagonal block. Test trials in the full block used all the classes shown in Fig. 1b (including the central class). Trials in the horizontal block only considered classes with *d*_2_ = 370 ms. Finally, trials in the diagonal block only used the diagonal classes (classes labeled from 1 to 10, Fig. 1b). Horizontal classes from the full block were used to compute psychometric curves (explained later). The horizontal and the diagonal blocks were employed to verify that the networks learned that these blocks define different contexts (Fig. 3d-e). All the other analyses were based on the diagonal block. Behavior was analyzed with 50,000 test trials per network. Analyses of neural activity were done with 4,000 test trials (when the sensory history was not considered) or with 20,000 test trials (for studies conditioning on the preceding class).

#### Training and simulations with several delay-period durations

In a second numerical experiment (Fig. 2b, left), fifteen networks were trained to perform the task with four different values of the delay period. In each training trial, one of the four possible durations was randomly selected from the set: 1,000 ms, 1,500 ms, 2,000 ms and 2,500 ms. Once a network was trained, a block of 4,000 consecutive test trials was generated for each duration of the delay-period, selecting classes from the diagonal block.

#### Simulations adding Gaussian noise in the neural activity

Once networks have been trained following the main numerical experiment, Gaussian noise with zero mean and standard deviation SD was applied to the membrane potential (Eq. (4)) either during the presentation of the first stimulus or during the delay period. From example network 1, two datasets of 50,000 (to study behavior; Fig. 2b, right and Extended Data Fig. 8) and 4,000 test trials (to analysis the neural activity; Extended Data Fig. 8) were generated per each considered value of SD (10 mV, 15 mV, 20 mV or 25 mV) and noise application stage (*d*_1_ or delay period).

#### Training and simulations with neural activity reset

In a third numerical experiment (Fig. 2c), 2,000 ms before the Start Cue of the following trial, the neural activity was reset to a random initial state, both on training and test trials. Seven networks were trained and tested with this experiment. A dataset of 4,000 test trials from the diagonal block and another dataset of 4,000 test trials from the full block were obtained for each network.

### Data analyses: Behavior

#### Psychometric function

This curve plots the probability *P*(*S*) to decide “*d*_1_ *< d*_2_” as a function of the values of first interval, *d*_1_ (Fig. 1d; Fig. 2c, left; Extended Data Fig. 1a; Extended Data Fig. 2a; Supplementary Fig. S2). It is computed using test trials with horizontal classes from the full block. It was fitted to a logistic sigmoid

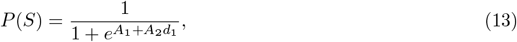

where *A*_1_ and *A*_2_ are adjustable parameters.

#### Discrimination function (accuracy)

This curve is the percentage of correct trials as a function of the class number (Fig. 1e; Fig. 2a and d, middle; Extended Data Fig. 1b and c; Extended Data Fig. 2b). It uses test trials with classes from the diagonal block. Its purpose is to analyze the network discrimination properties.

#### Phenomenological measure of the contraction bias

Denoting the probability of deciding *S* given a presented class *C* as *P*(*S*|*C*), a phenomenological measure *B* of the contraction bias (Fig. 1g; Fig. 2c, right; Fig. 5c and f; Fig. 6) was defined as the root-mean-square deviation of *P*(*S*|*C*) on diagonal classes, relative to the classes with *d*_1_ = 370 ms (classes 3 and 8 in Fig. 1b)

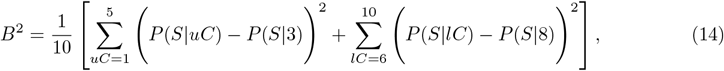

where *uC* and *lC* denote classes on the upper and the lower diagonals of the stimulus set (Fig. 1b), respectively. Note that the value of *B*^2^ corresponding to the maximum possible bias equals 2*/*5, while for the minimum possible bias it is 0.

### Data analyses: Neural Activity

#### Firing Rates

Trials were aligned at the relevant task events: SO1 (presentation of the first stimulus), SF1 (delay period) or SF2 (inter-trial-interval, ITI). Firing rates were calculated by sliding a 150 ms window by 25 ms intervals, except in the analysis of the relationship between behavior and activity (see below), where a time window of size 100 ms sliding every 10 ms was used. Trial-averaged firing rates were estimated conditioning over the value of *d*_1_.

#### Neural Tuning

During the delay period, the firing rates of a large population of neurons coded the duration of the first time interval parametrically, either in a monotonically increasing or decreasing fashion. For this classification, we fitted the firing rate of each neuron with a linear function *αd*_1_ + *β*. If the slope of the fit was significant and positive, then the neuron was said to be positively tuned and it was classified as *plus*. Conversely, if the slope was significant and negative, the neuron was negatively tuned and it was classified as *minus*. From the 500 neurons of the example network 1, 137 neurons were classified as plus and 97 as minus, using test trials from the diagonal block. In the full block there were 135 plus neurons and 90 minus neurons. Example network 2 had 116 plus neurons and 114 minus neurons using the diagonal block while with the full block there were 115 plus neurons and 97 minus neurons.

#### Mutual Information

The information that neurons carried about the duration of the intervals was measured by the mutual information (MI) [60]. The single neuron response was quantified as a discrete array *r* = {*r*_1_, …, *r*_*L*_}, where each *r*_*t*_ indicates the number of spikes fired in the *t*-th time bin and *L* is the total number of bins. The bins, of size Δ*t* = 250 ms, were shifted by steps of 25 ms. *D*_1_ collected all possible values of *d*_1_ and *R* collected all values of *r*. The MI about a time interval *d*_1_ carried by a neuron is

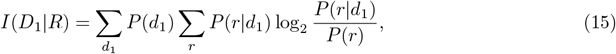

where *P*(*d*_1_) is the probability of a time interval *d*_1_ and *P*(*r*|*d*_1_) is the conditional probability of observing a response *r* given that the presented interval was *d*_1_. To compute Eq. (15) we used a MATLAB-based toolbox [61] that includes corrections for a bias in the estimate of the MI due to limited-sampling of the neural response [62]. This bias was estimated using an extrapolation quadratic in the number of trials, *N*_*trial*_,

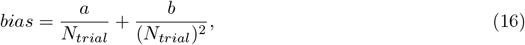

where *a* and *b* are free parameters.

Once the MI from each single neuron was obtained (Supplementary Fig. S3), we analysed its significance. For that, we used a bootstrap procedure given by the toolbox, which consists in pairing randomly stimuli and responses in order to destroy its correlation [61]. The MI of a single neuron at a given time was significant if its value was higher than the sum of the corresponding bootstrapped MI and the standard deviation over permutations used in the bootstrap procedure. Then, the percentage of neurons with significant MI through time was calculated (Fig. 4c-d). We implemented a second shuffling procedure to destroy causality between the responses of consecutive trials. This shuffling consisted of putting together all firing rates of all trials of all neurons and performing 25 random permutations without replacement. From this shuffled dataset, we calculated, bin-by-bin, the percentage of neurons with significant mutual information and its mean ± CI’s (grey horizontal shadings in Fig. 4c-d).

### Data analyses: State-space Analysis and Geometry

#### *Principal Component Analysis*(PCA)

We used PCA to examine the trial-averaged firing rates and to identify the low-dimensional space that captured at least 90% of data variance. Neural activity described curved trajectories embedded in this low-dimensional space; the associated speed was computed as the norm of the covered distance by a time window of 150 ms, which slides 25 ms (Fig. 3c).

#### KiNeT Method for relative distances

Relative distance between a pair of trajectories (*i* and *j*) was defined following the Kinematic analysis of Neural Trajectory (KiNeT) methodology [19]. We first selected one of the two trajectories as a reference (e.g., *ref* =*i*) and then defined corresponding states between the reference and the other trajectory (*j*) as follows: If at time *t* after the aligning event (SO1 for the presentation epoch of *d*_1_, SF1 for the delay period and SF2 for the ITI) the reference trajectory was in the state *s*_*ref*_ (*t*), then the corresponding state *s*_*j*_(*t*) on trajectory *j* was the one at the minimum Euclidean distance to *s*_*ref*_ (*t*). This is the relative distance *D*_*j*_(*t*) between the two trajectories. Relative distances were computed in a PC space which accounted for at least 90% of the total variance during the investigated epoch. This analysis has been used in the following situations:

1. State-space trajectories depend on context. To verify that classes common to horizontal and diagonal blocks (classes 5 and 6) defined different trajectories (Fig. 3d-e), their relative distances were computed, taking the trajectory in the diagonal context as a reference. Standard errors were computed using the mean ± 95% confidence intervals (CI), calculated from the PCA and the corresponding KiNeT analysis performed on each of the 100 bootstrap resamples with replacement.
2. State-space trajectories depend on *d*_1_. As a first check whether the duration of the first interval was transmitted to the delay interval and the ITI, we verified that the relative distances between trajectories with different values of *d*_1_ were significantly nonzero (Fig. 4a-b). Standard errors were computed as in the previous situation.
3. Relative distances have been also used to examine whether they encoded an estimator of *d*_1_ during the delay period (Fig. 7d; Extended Data Fig. 7d; Extended Data Fig. 9g and j). See details below.

### Bayesian Model

#### Formulation and Likelihood

Behavioral data were fitted using a Bayesian model. We assumed that at the current trial *n* the Bayesian observer makes an observation of the first interval (*o*_1,*n*_) given by a combination of the current-trial noisy measurement of *d*_1_ 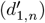 and the observation in the preceding trial

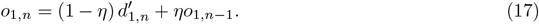

Here *η* is a parameter that measures the contribution of the observation in the preceding trial (if *η* = 0 there is no contribution from sensory history while if *η* = 1 the current stimulus does not contribute). To work with analytical expressions, instead of iterating this equation we expressed *o*_1,*n*_ by extracting the interval measurements made in the *m* past trials, named as the short-term sensory history (Eq. (1) in the main text)

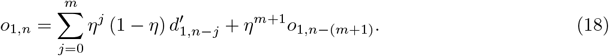

We assumed that the noisy measurements of the intervals can be described by normal distributions (*j* = 0, 1, …, *m*)

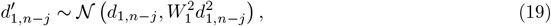

where *W*_1_ is the Weber fraction, a parameter of the Bayesian model to be determined. In addition, we assumed that trials prior to the last *m* constitute a Gaussian stationary regime (the long-term sensory history). The mean and variance of the stationary variable are obtained by self-consistency arguments

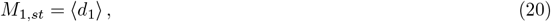

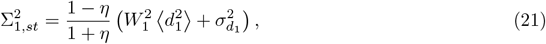

where ⟨*d*_1_⟩ and 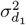 are the mean and variance of the prior probability of *d*_1_. Within this Gaussian approximation the observation in the current trial is also a Gaussian variable, with mean ℳ and variance 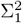, that we denote by 𝒪_1_

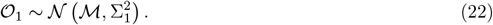

The distribution of this variable defines the likelihood of an observation 𝒪_1_, given the short-term sensory history, that we denote by *P*(𝒪_1_|*d*_1,*n*_, …, *d*_1,*n−m*_). The mean and variance of the observation probability for the general case, in which the transient consists of the previous *m* trials, is given by

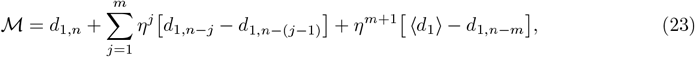

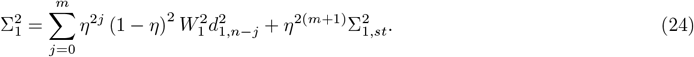

We considered two cases of the model, *m* = 0 and *m* = 1. In the first case all past trials contribute to the stationary regime. The likelihood *P*(𝒪_1_|*d*_1,*n*_) has mean and variance

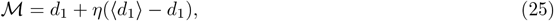

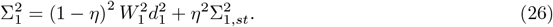

In the second case (*m* = 1) the transient regime contains only the preceding trial. The likelihood *P*(𝒪_1_|*d*_1,*n*_, *d*_1,*n−*1_) has mean and variance

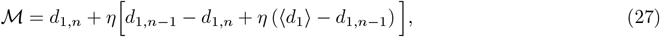

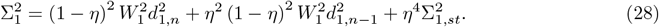

A more general model, which considers the contribution of the noisy measurements of the second interval in the previous trial 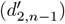, is given in Supplementary Text S1.

#### Beliefs

The computation of posterior probabilities closely follows a Bayesian model for tactile frequency discrimination [9]. For the sake of completeness, here we adapt it to the time interval discrimination task. The main difference is that the likelihoods are given by the expressions derived above. We present the equations only for the case *m* = 1. The discrete prior probabilities of the first interval are denoted as 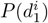 (*i* = 1, …, 5 labels the possible values of *d*_1_ in the stimulus set, Fig. 1b). Bayes theorem establishes that the posterior distribution 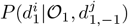, conditioned to the preceding interval 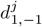 (*j* = 1, …, 5), results from combining the noisy observation with the prior distribution,

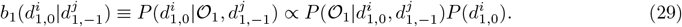

Notice that the notation was simplified denoting *d*_1,*n*_ and *d*_1,*n−*1_ as *d*_1,0_ and *d*_1,*−*1_, respectively. The left hand side indicates that this probability is also known as the belief 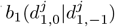 about the duration of the first time interval, conditioned to the duration in the previous trial. One may need to marginalize over 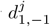, summing over all values of *j*.

Denoting the discrete prior probabilities of the second time interval as 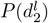 (*l* = 1, …, 9 labels the possible values of *d*_2_ in the stimulus set, Fig. 1b), the posterior probability 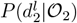, or belief 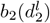 about the duration of the second time interval, is

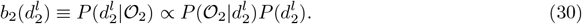

Let us denote class *k* as 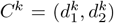, where *k* = 1, …, 10 is a class label (Fig. 1b). The joint posterior probabilities (or joint beliefs) that class *C*^*k*^ had been presented in the current trial (the observations are 𝒪_1_ and 𝒪_2_), conditioned to the interval in the previous trial are [9]

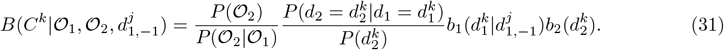

The first quotient is a normalization factor. 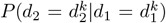 is the transition probability matrix of *d*_2_ being presented after *d*_1_. If the network has good knowledge of the structure of the set, the only non-zero transition matrix elements correspond to the 10 classes in the set. Note that this matrix depends on the block under consideration (full, diagonal or horizontal blocks, Fig. 1b). For example, in the full block, for a given *d*_1_ the second interval can take either one or two or three different values. Thus, the non-zero transition probabilities would be 1, 1/2 or 1/3, respectively (Fig. 1c).

The Bayesian observer makes a decision based on which of the choices, “*d*_1_ *< d*_2_” or “*d*_1_ *> d*_2_”, has a higher belief. These beliefs are computed by adding the class beliefs *B*(*C*^*k*^|𝒪_1_, 𝒪_2_) with a specific choice

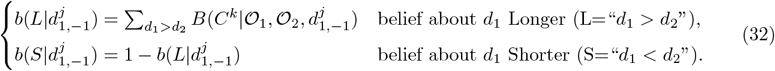

#### Probability of deciding “d_1_ < d_2_”

For the model version *m* = 0, the probability that the Bayesian observer, with knowledge of 𝒪_1_ and 𝒪_2_, chooses “*f*_1_ *< f*_2_” when the current class is 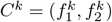 is

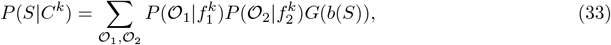

where *G*(*b*(*S*)) = 1 if *b*(*S*) *>* 0.5 and zero otherwise [9], [15].

Likewise, for the model version *m* = 1, the conditional probability to choose “*d*_1_ *< d*_2_” when, in addition, the previous class is 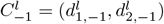 is

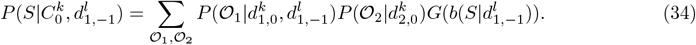

#### Choice bias

The trained networks showed a tendency to repeat the choice of the previous trial when it had been correct or make the opposite choice when it had not, that is, they followed a win-stay/loss-shift (WSLS) strategy. We modelled this choice bias by modifying the argument of the decision factor *G* in Eq. (34) which now depends on the outcome of the previous trial, *U*_*−*1_, that takes the values 1 or 0 for correct and incorrect previous choices, respectively. Moreover, the choice bias depended on which diagonal of the stimulus set (Fig. 1b) the previous class was on. So, instead of Eq. (34) we now have

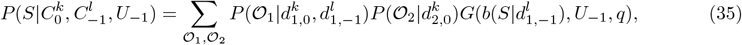

where *q* = 1 (*q* = 2) denotes the upper (lower) diagonal. Now the function *G* is

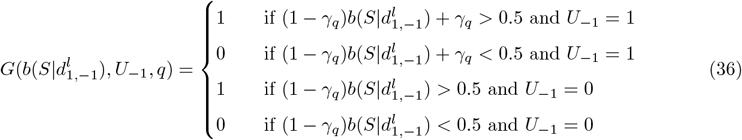

where the parameters *γ*_1_, *γ*_2_ bias choices *S* and *L* according to the WSLS strategy. Notice that Eq. (35) yields 20 performance curves defined on current classes (one per previous class and outcome), which can be obtained from the simulated behavioral data (Fig. 2a).

#### Correlations between the bias B and the model parameters

Correlations (between the model parameters with each other and with the bias *B*) in Fig. 6 were computed with the MATLAB function *corrcoef*. Correlations were considered significant if the p-value was smaller than 0.05.

### Fitting the Bayesian Model to Data

Both versions of the model (*m* = 0 and *m* = 1) were fitted by minimizing mean squared differences between the model and network performances. Minimizations were done using MATLAB’s *simulannealbnd* function which implements a simulated annealing algorithm. For the case *m* = 0, the model fit (Fig. 5a) was done by minimizing the mean squared difference between the performance curve in Eq. (33) and the corresponding behavioral curve obtained from simulations. For the case *m* = 1 (conditioning over the previous class), the model fit was done using the 20 performance curves in Eq. (35). The minimized error resulted from summing over the 20 squared differences between the model and the network performances (one difference per previous class and outcome). Then, from the obtained fit we computed the 10 accuracy curves conditioned to the previous class (Fig. 5d) and the marginal accuracy (Fig. 5g), by marginalization, as explained next.

#### Marginalization over previous class and outcome

From the simulated behavioral data we know the performance curves 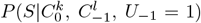, conditioned on the current class 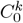, the previous class 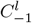 and the previous outcome *U*_*−*1_. From these curves, the parameters of the version *m* = 1 of the model were obtained as described above. Now, we want to obtain the fitted marginal discrimination curve 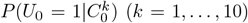 by marginalizing 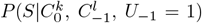 on the previous class and outcome. For this, it is necessary to solve a system of 10 equations with 10 unknowns (for further details see Supplementary Text S2). The first 5 equations of this system correspond to the 5 current classes on the upper diagonal (*k* = 1, …, 5)

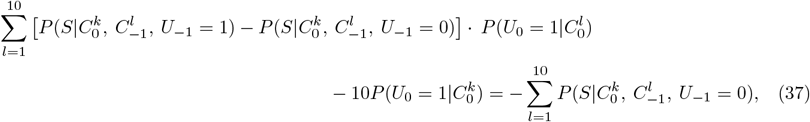

with 10 unknowns 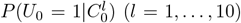 (*l* = 1, …, 10). The other 5 equations correspond to the 5 current classes on the bottom diagonal (*k* = 6, …, 10)

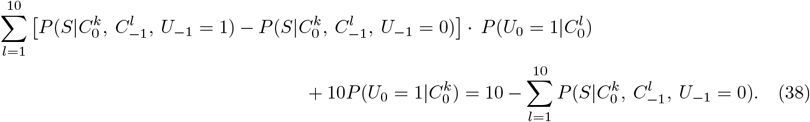

Solving the system in Eqs. (37-38) we obtained the marginal accuracy curve, 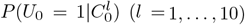 (Fig. 5g). Once this is known, the probability of deciding *S* ≡ “*d*_1_ *< d*_2_” conditioned on the current and previous classes, *P*(*S*|*C*_0_, *C*_*−*1_) (Fig. 5d), can be computed from

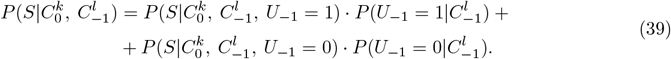

### Model comparison

#### *Akaike Information Criteria*(AIC)

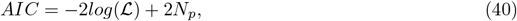

where *N*_*p*_ is the number of model parameters and ℒ is the likelihood of obtaining the observed choice (“*d*_1_ *< d*_2_” or “*d*_1_ *> d*_2_”) across all trials. For given class and choice, the likelihood function takes the values *l*(*x* = “*d*_1_ *< d*_2_”) = *P*(“*d*_1_ *< d*_2_”) or *l*(*x* = “*d*_1_ *> d*_2_”) = *P*(“*d*_1_ *> d*_2_”). Then, *log*(ℒ) is the sum over all trials

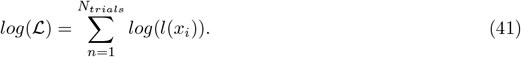

#### *Bayesian Information Criteria*(BIC)

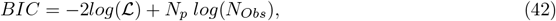

where *N*_*Obs*_ is the sample size (number of observations).

### Estimation of *d*_1_ and its relationship with state-space geometry

To check whether geometric features of the trajectories encoded an estimate of the first interval we compared separately three estimators (the true value of *d*_1_, the Bayesian estimator *d*_1,*Bayes*_ and the ML estimator *d*_1,*ML*_) with measures of the curvature, relative distances and spatial structure of the quasi-attractor. To do this, we first computed the estimators in single trials simulating the model (version *m* = 1, Eqs. (27,28)). Then, we obtained the means *d*_1,*Bayes*_ and *d*_1,*ML*_ as an average over trials.

#### Bayesian and Maximum Likelihood estimators

For a given trial *n*, the Bayesian estimator of *d*_1,*n*_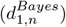 results from a loss function; taking it as the squared error

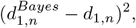

the Bayesian estimator is the mean of *d*_1_ weighted with the posterior probability *b*_1_ 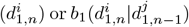

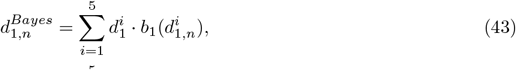

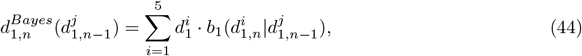

for the marginal and conditional cases, respectively, from model version *m* = 1. The ML estimator 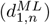 is the value of *d*_1_ that maximizes the likelihood function,

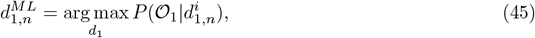

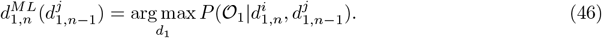

Once the model parameters have been determined, we simulated 10,000 trials (marginal case) and 10,000 trials for each 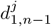 (conditional case) and computed the corresponding Bayesian (Eqs. (43-44)) and ML (Eqs. (45-46)) estimators. Their trial averages yielded *d*_1,*Bayes*_ and *d*_1,*ML*_, respectively. The bias of the Bayesian estimator (Supplementary Fig. S8) was obtained from

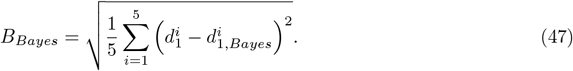

#### Curvature of the one-dimensional curved manifold

To investigate the state-space representation of *d*_1_ during its presentation epoch we analyzed the curvature of the one-dimensional manifold developed during that period [21]. We first projected the neural trajectories onto an encoding axis, **u**, defined as a unit vector between the state *s*(t=SF1_1_) associated to the shortest SF1, and the state *s*(t=SF1_5_) associated to the longest SF1. Then, for each trajectory *j*(*j* = 1, …, 5) we computed the projection, *Proj*_*j*_, of the state *s*(t=SF1_*j*_) onto the encoding axis

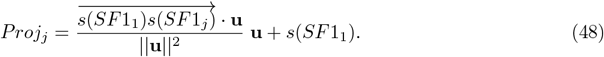

To test which of the three estimators best explained the encoding axis projection data during the presentation of *d*_1_ we fitted them, separately, to

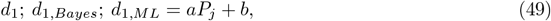

where *P*_*j*_ (*j* = 1, …, 5) are the distances between projections on an encoding axis [21], taking as reference the projection of the shortest *d*_1_. Again, *d*_1,*Bayes*_ and *d*_1,*ML*_ are the average of the estimators over the simulated trials obtained either with Eqs. (43) and (45) (marginalizing over the previous stimulus) or with Eqs. (44) and (46) (conditioning over the previous stimulus). Firing rates were computed using a 100 ms analysis window, with a slide of 10 ms throughout the presentation of the first interval. We kept the number of PCs needed to explain, at least, 95% of the variance (between 3 and 5 PCs). This analysis was done for the marginal case (Fig. 7a-c and Extended Data Fig. 7a-c) and the conditional case (Extended Data Fig. 9a-f).

#### Mean relative distances during the delay period

To test which of the three estimators best explained the relative distances between neural trajectories (taking as reference the trajectory corresponding to *d*_1_=370 ms; KiNeT method) during the delay period, we compared the three hypotheses

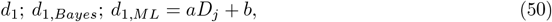

where *D*_*j*_ (*j* = 1, …, 5) is the temporal mean of the relative distances [20]. We used an analysis window of 100 ms with a slide of 10 ms throughout the delay period. We kept the number of PCs needed to explain, at least, 90% of the variance (between 5 and 11 PCs). This analysis was done for the marginal case (Fig. 7d-f and Extended Data Fig. 7d-f) and the conditional case (Extended Data Fig. 9g-l).

#### Distances between terminal states on the quasi-attractor

Likewise, to check which estimator is encoded by the attractor we considered the Euclidean distances *δ*_*j*_ (*j* = 1, …, 5) between the terminal states on the attractor region of each *d*_1_ trajectory and the shortest *d*_1_ trajectory.

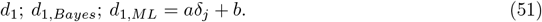

The test was done in the last 100 ms of the delay epoch. Although for network 1 the first PC explained about 96% of the variance, the quality of the regressions improved by including the second PC (which contributed 3.5% of the variance). This analysis was done for the marginal case (Fig. 7g-i and Extended Data Fig. 7g-i) and the conditional case (Extended Data Fig. 9m-r).

In these three tests the goodness of the fits was compared using their Root Mean Square Errors (RMSEs). Significance was evaluated using a bootstrap procedure. From the original neural population we created 100 resamples by repeatedly sampling with replacement. Then we repeated the tests in each resample and computed the corresponding dispersions.

### Compression of geometric features

Denoting by *Z*_*j*_ (*j* = 1, …, 5) any of the three geometric features considered in this work (projections on the encoding axis during presentation of *d*_1_, *P*_*j*_; mean relative distances during the delay period, *D*_*j*_ = ⟨*D*_*j*_(*t*)⟩; distances to terminal states on the attractor region, *δ*_*j*_), we defined the compression *C*[*Z*] as

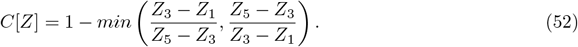

## Code and Data Availability

Training with full-FORCE was done adapting to the time interval discrimination task the MATLAB codes from [36], available in https://github.com/briandepasquale/supervised_learning_recurrent_spiking_networks.

Custom MATLAB scripts used to implement the model and produce the figures will be available upon publication. The simulated data that support the findings in this study are available from the corresponding author upon reasonable request.

## Acknowledgements

This work was supported by the grant PGC2018-101992-B-I00 from the Spanish Ministry of Science, Innovation and Universities. We thank Joan Falcó-Roget and Yonatan Loewenstein for discussions.

## Author contributions

L.S.-F. and N.P. designed the study. L.S.-F. implemented the training methods and performed the numerical analysis. L.S.-F. and N.P. developed the analysis techniques. M.B. and N.P. designed the normative model. L.S.-F., M.B. and N.P. performed research and contributed to writing the manuscript.

## Extended Data Figures

**Extended Data Fig. 1.**
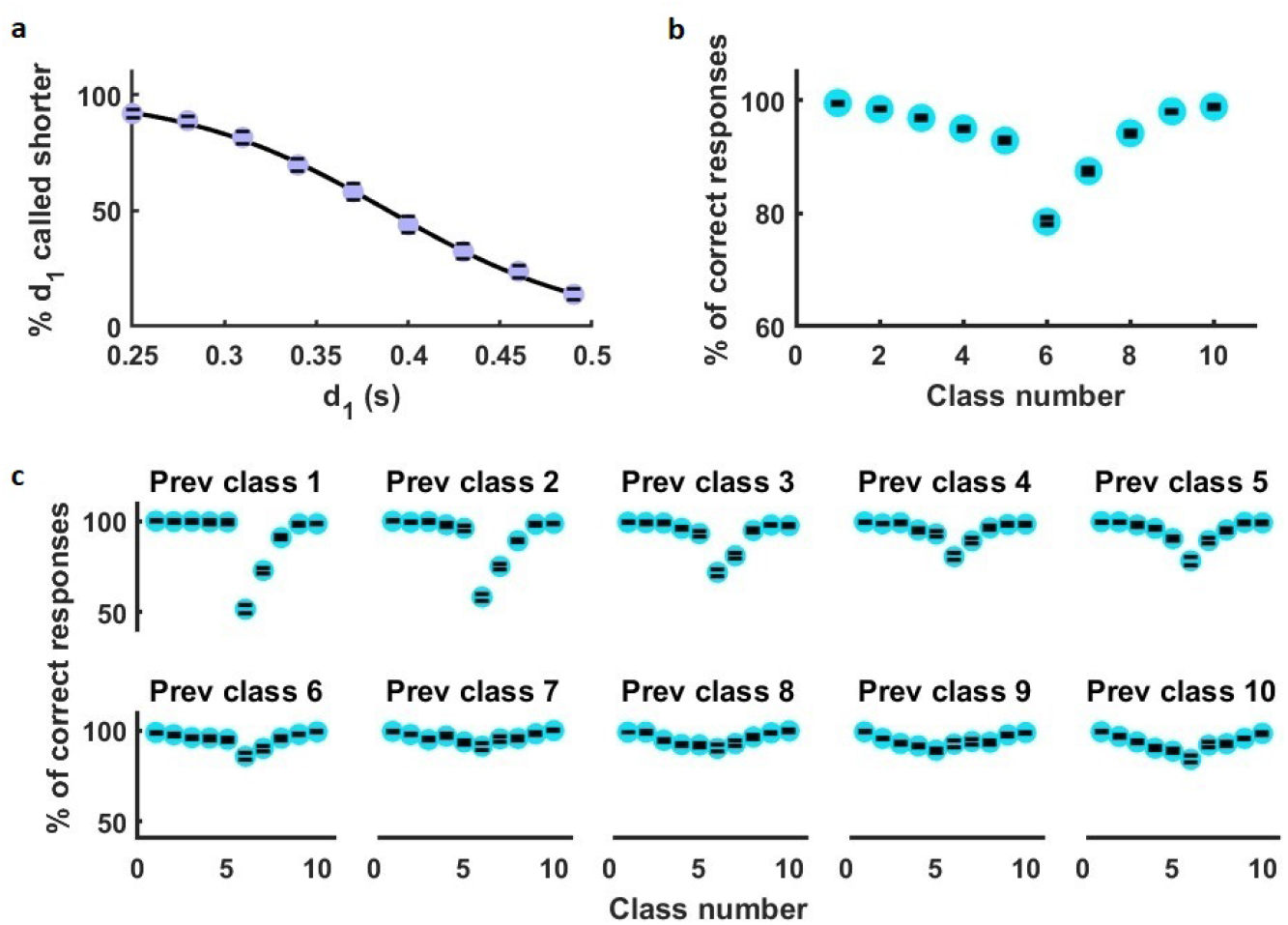
Simulated behavioral data of example network 2. **a**. Psychometric curve (using horizontal classes from the full block). Black line is a sigmoidal fit of the simulated data. **b**. The accuracy curve shows the percentage of correct test trials (using the classes from the diagonal block). **c**. Ten accuracy curves, each conditioned over one of the ten possible previous classes of the diagonal block. Error bars were computed with 1,000 bootstrap resamples.

**Extended Data Fig. 2.**
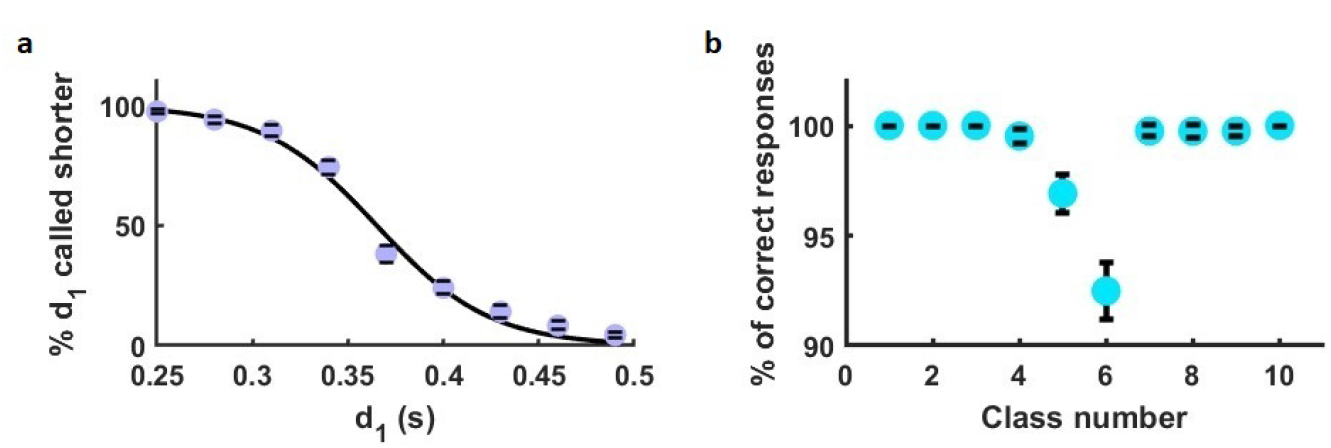
Example network trained and tested resetting its neural activity. **a-b**. Simulated behavioral data. The reset took place 2,000 ms before the end of each training and test trial. **a**. Psychometric curve. Black line is a sigmoidal fit. **b**. Accuracy curve. Error bars were computed with 1,000 bootstrap resamples.

**Extended Data Fig. 3.**
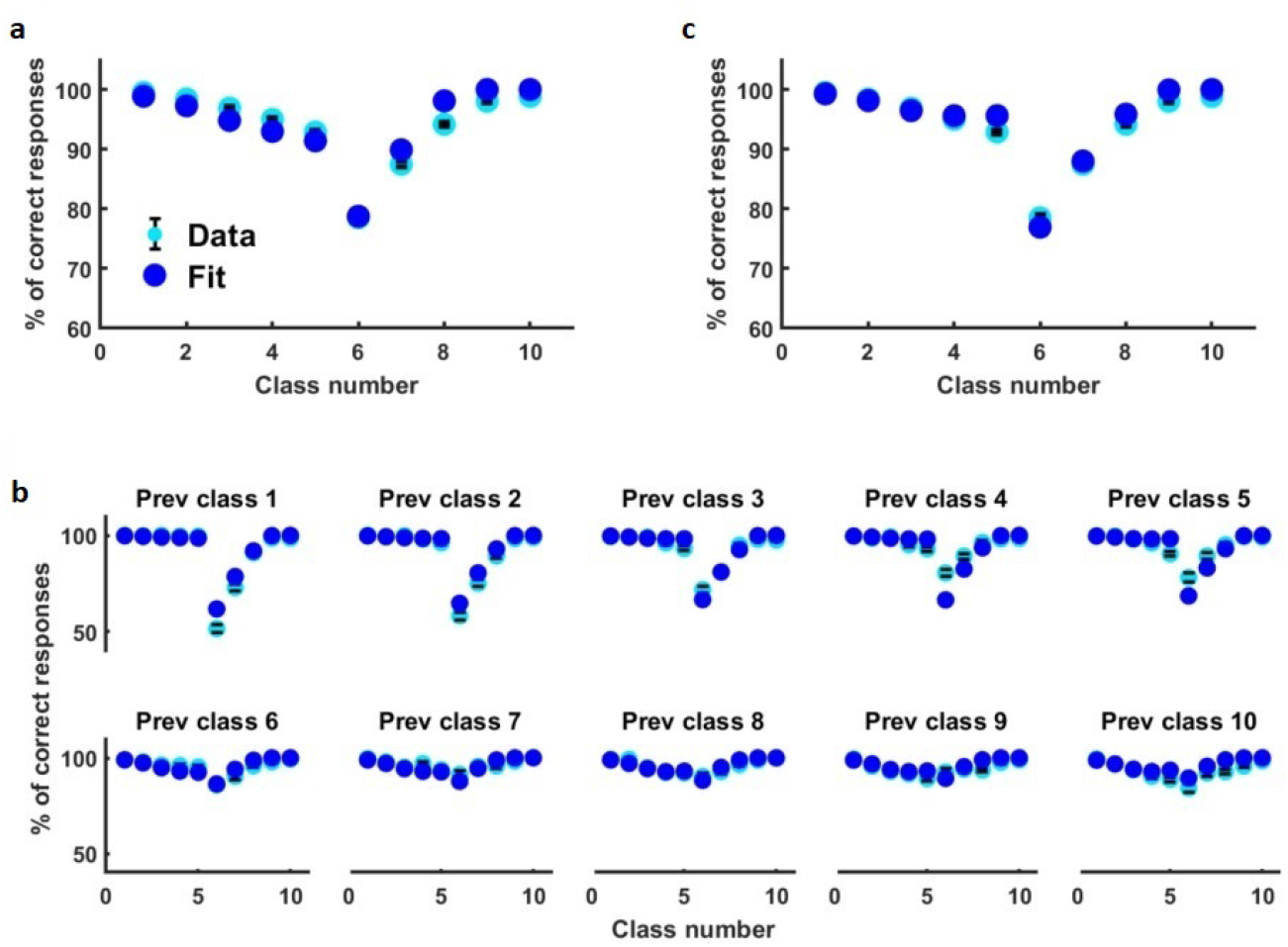
Fits of the behavioral data of example network 2. **a**. Fit of the accuracy curve with the model version *m* = 0 (Eqs. (2a,b)). Cyan circles with error bars are the simulated behavioral data of example network 2 in Extended Data Fig. 1b. Dark blue circles are the fit. **b**. Fit of the ten accuracy curves conditioned over each possible preceding diagonal class, obtained with model version *m* = 1 (Eqs. (3a,b)). Cyan circles are the behavioral data of example network 2 in Extended Data Fig. 1c. **c**. Marginal accuracy curve obtained from the fit in (b) by marginalizing over both the class and the outcome in the previous trial (see Methods).

**Extended Data Fig. 4.**
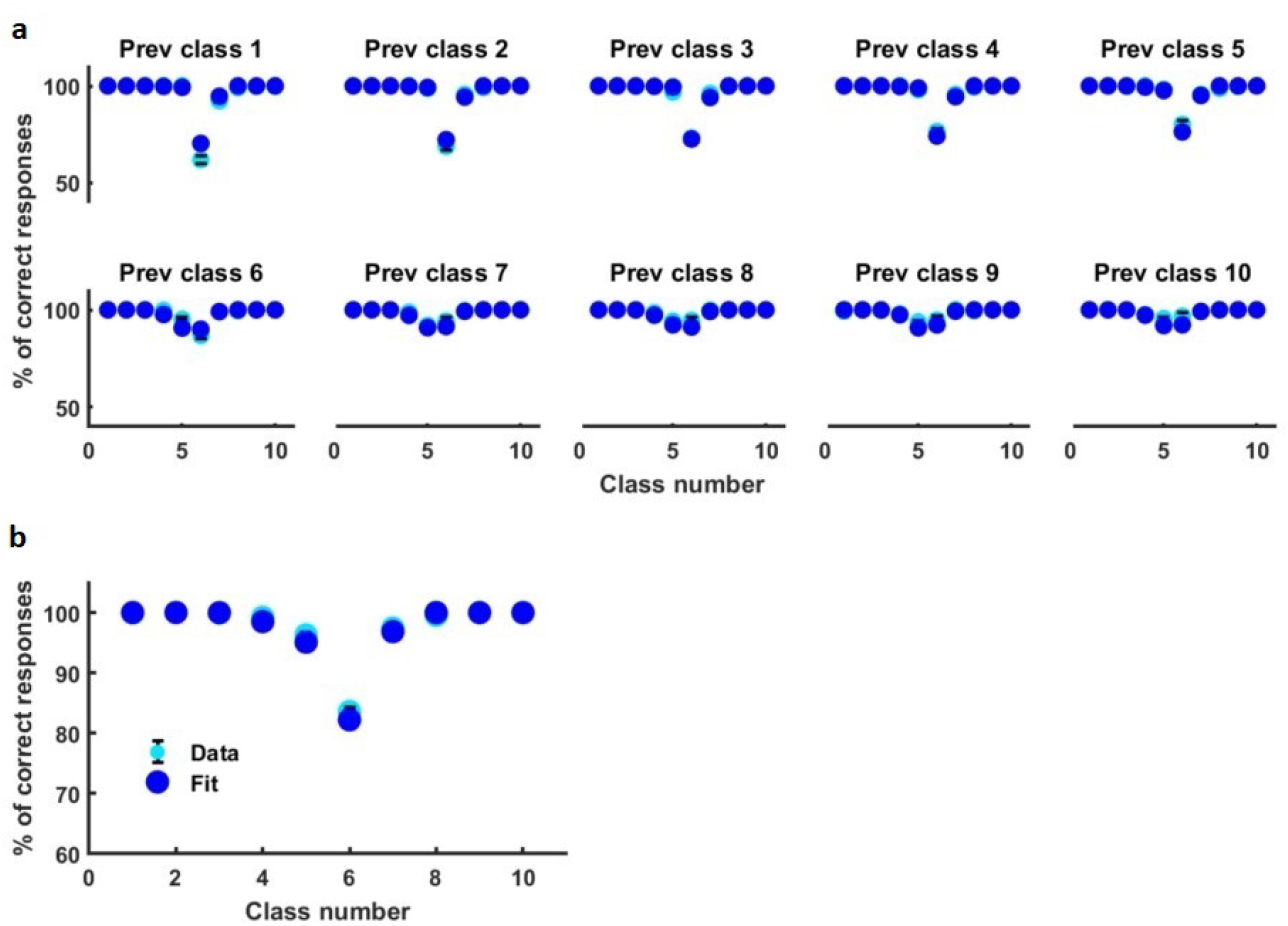
Model that includes both the previous first and second intervals. **a-b**. Fit of the behavioral data of network 1. **a**. Fit of the ten accuracy curves conditioned over each possible preceding diagonal class (Eqs. (S10, S11)). Cyan dots with error bars denote the behavioral data of example network 1 in Fig. 2a. Dark blue circles are the fit. The model is described in Supplementary Text S1. **b**. Marginal accuracy curve, obtained by marginalizing the fitted curves over both the class and the outcome in the previous trial (see Methods). Cyan circles are the behavioral data of example network 1 in Fig. 1e, right.

**Extended Data Fig. 5.**
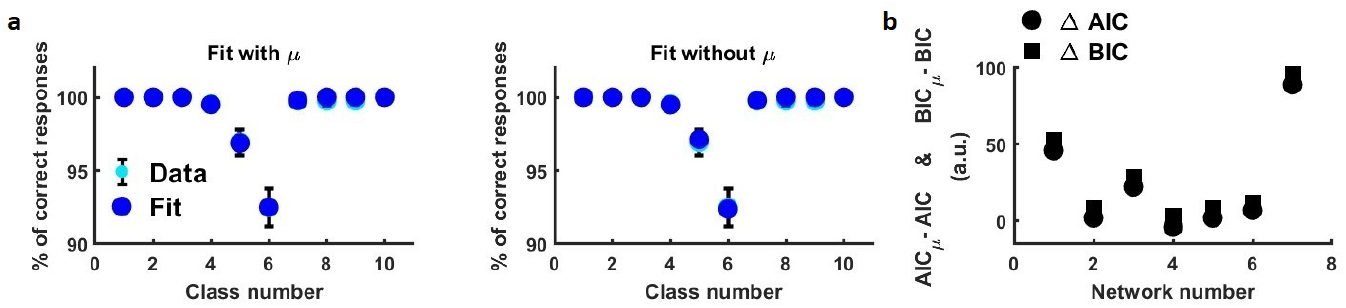
Evaluation of the hypothesis that a reset network has knowledge about the statistics of *d*_1_. **a**. Two fits of simulated behavioral data of an example network trained and tested resetting its neural activity (Extended Data Fig. 2b). **Left**: Fit with a model that depends on the mean duration ⟨*d*_1_⟩ (*μ* ≠ 0). **Right**: Fit with a model that does not depend on it (*μ* = 0). **b**. Difference between the AIC measure for the model with parameter *μ* and the AIC for the model without this parameter, versus network number (circles). Id. for the BIC measures (squares). Comparison of the two models tends to favor the one that does not require knowledge of ⟨*d*_1_⟩.

**Extended Data Fig. 6.**
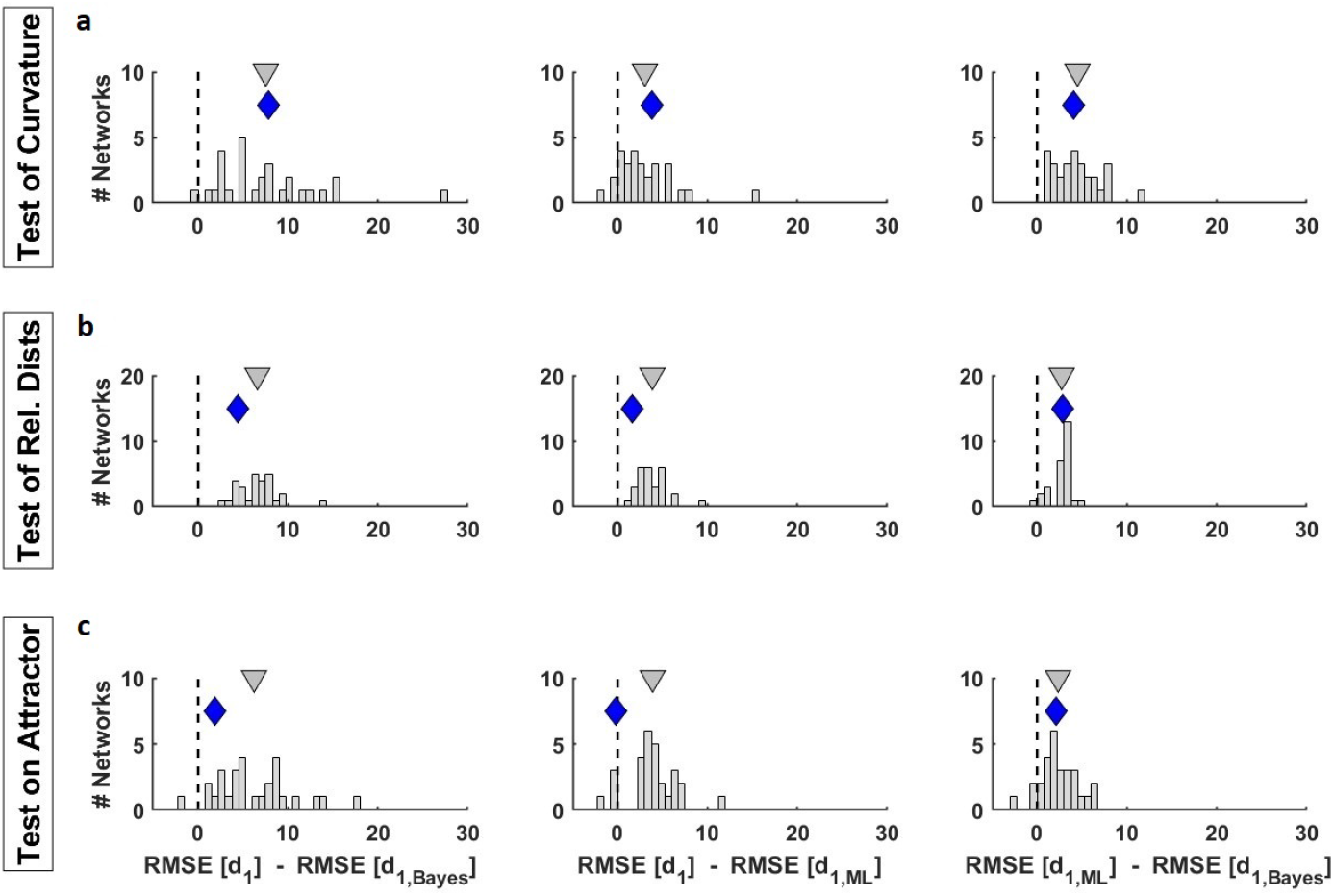
Geometric features best encode the Bayesian estimator in all positive networks (model version *m* = 1). Histograms over networks of RMSE differences resulting from (**a**) regressions of distances between projections, *aP*_*j*_ + *b*, (**b**) regressions of temporal means of relative distances, *aD*_*j*_ + *b* and (**c**) regressions of distances between terminal states on the attractor, *aδ*_*j*_ + *b*. The mean RMSE differences between the Bayesian estimator and the true value of *d*_1_ (left) was around two times larger than that of the ML estimator (middle). In the curvature test only one network favored the true *d*_1_ instead of the Bayesian estimator. Only one network preferred the ML estimator in the test of relative distances and only four positive networks did not choose the Bayesian estimator in the test on the attractor. This indicates that the hypothesis that geometric features (curvature, relative distances or distances on the attractor) encode the Bayesian estimator is better than the hypothesis that it encodes the ML estimator. Gray triangles indicate histogram means. Blue diamonds denote example network 1. Dashed vertical lines identify zero RMSE differences.

**Extended Data Fig. 7.**
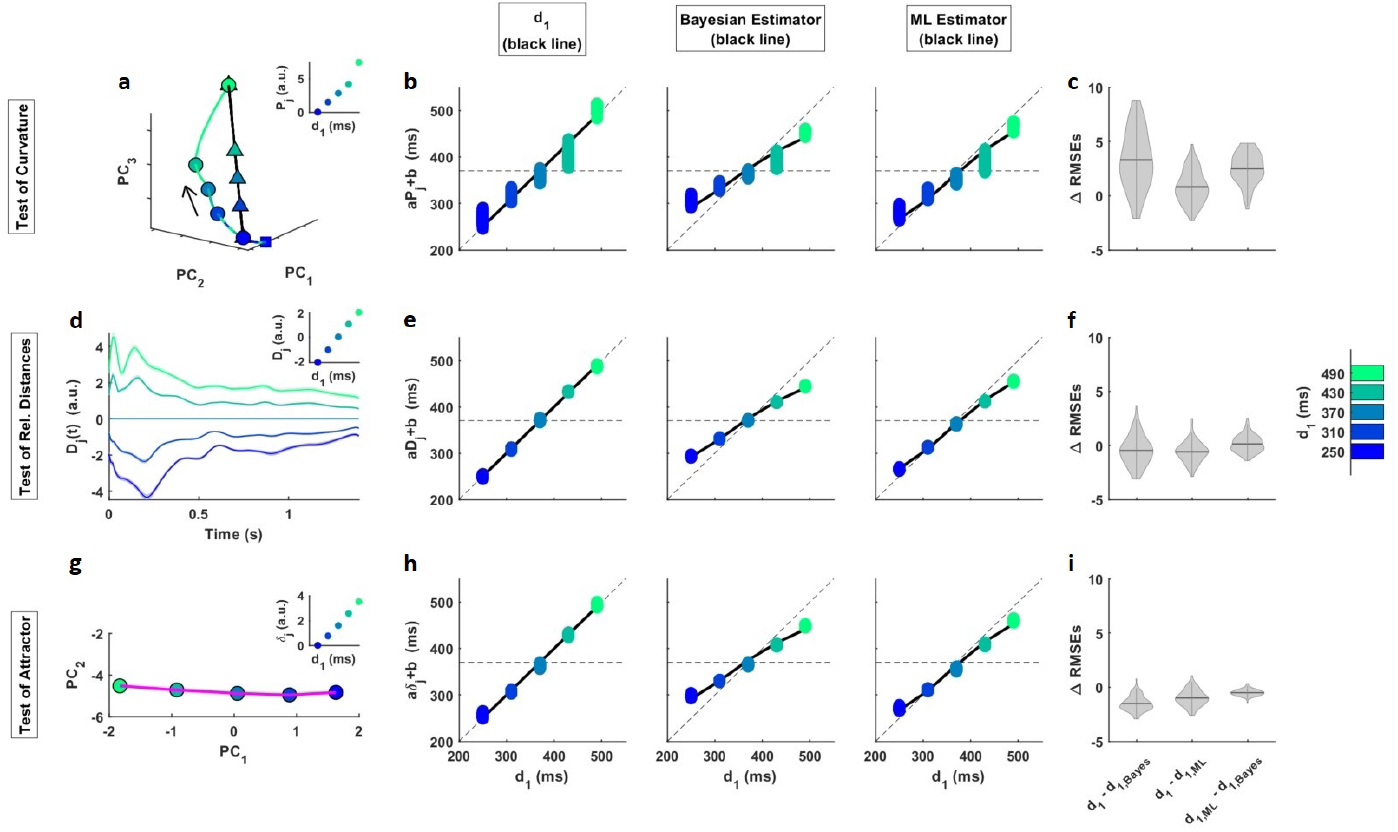
Several geometric features code the Bayesian estimator of *d*_1_. **a-i** Example network 2. **a-c**. Curvature of the one-dimensional curved manifold during the presentation of the first stimulus. **a**. Neural trajectories and projections (triangles) of each SF1 terminal state (circles) onto the encoding axis, **u** (black line). Squares represent the beginning of the figure, located within the first stimulus. Neural trajectories are embedded within a state space built with the first 4 PCs, which explain 95% of variance. **Inset** shows, as a function of *d*_1_, the Euclidean distances between projections, *P*_*j*_, (where *j* = 1, …, 5 labels the possible values of *d*_1_ in the diagonal block) taken as reference the projection of the shortest *d*_1_. **b**. Linear regression of the projections (*aP*_*j*_ + *b*) to the true *d*_1_ (left), the Bayesian estimator (middle) and the ML estimator (right), as a function of *d*_1_. Colored dots show multiple estimates of the regression derived from bootstrapping (*n*=100). Solid lines indicate the true *d*_1_ (left), the Bayesian estimator (middle) and the ML estimator (right) versus *d*_1_. **c**. Violin plots (kernel probability density) on shufflings of the RMSE differences between the three hypotheses (*d*_1_, Bayesian estimator and ML estimator). The width of the gray area indicates the density of the shufflings located there. Horizontal line inside of each violin plot indicates the distribution mean. The mean of the difference 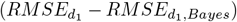 was approximately 2.5 times larger for network 1 than for network 2. **d-f**. Mean relative distances during the delay period. **d**. Relative distance, *D*_*j*_(*t*), between each trajectory *j* = 1, …, 5 and the reference trajectory *d*_1_=370 ms. Neural trajectories are embedded in a state space built with 7 PCs, which explain 91% of variance. Shadings indicate the mean ± 95% CIs computed from 100 bootstrap resamples. **Inset** shows the temporal means of relative distances as a function of *d*_1_. **e**. Linear regression of the mean relative distances (*aD*_*j*_ +*b*) to the true *d*_1_ (left), the Bayesian estimator (middle) and the ML estimator (right) as a function of *d*_1_ (same format as in (b)). **f**. Analogous to (c), but for the relative distance analysis. Two thirds of the resamples prefer the real *d*_1_ hypothesis. **g-i**. Distances between terminal states on the quasi-attractor. **g**. One-dimensional quasi-attractor region (magenta line), at the end of the delay period, embedded within the first 2 PCs space (99.6% of variance). **Inset** shows the Euclidean distances, *δ*_*j*_, between two *d*_1_ terminal states (taken the shortest *d*_1_ as reference) on the attractor region, as function of *d*_1_. **h**. Linear regression of the terminal states on attractor (*aδ*_*j*_ + *b*) to the true *d*_1_ (left), the Bayesian estimator (middle) and the ML estimator (right) as a function of *d*_1_ (same format as in (b)). **i**. Analogous to (c), but for the terminal states analysis. The attractor was analyzed with a time window of size 100 ms at the end of the delay period. In the vast majority of shufflings, the real *d*_1_ hypothesis is chosen.

**Extended Data Fig. 8.**
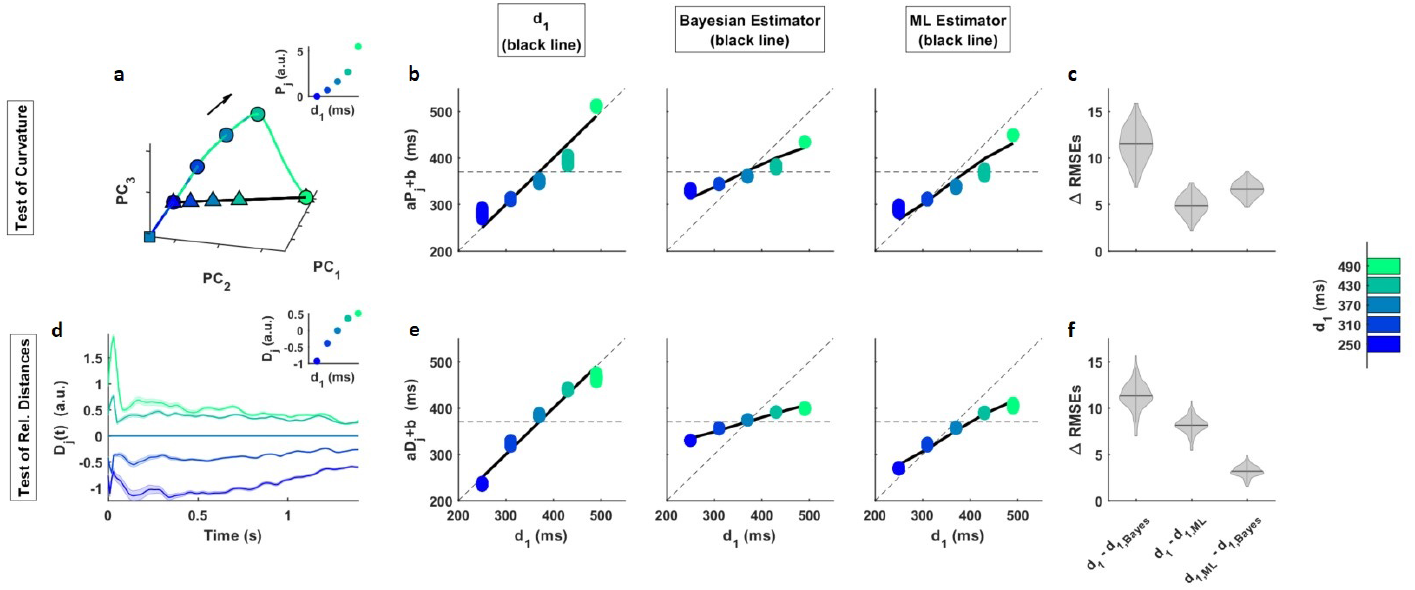
Gaussian noise increases the compression of geometric features of state space trajectories, which still code the Bayesian estimator of *d*_1_. **a-f** Example network 1. **a-c**. Curvature of the one-dimensional curved manifold during the presentation of the first stimulus perturbed by Gaussian noise with SD = 25 mV. **a**. Neural trajectories and projections (triangles) of each SF1 terminal state (circles) onto the encoding axis, **u** (black line). Squares represent the beginning of the figure, located within the first stimulus. Neural trajectories are embedded within a state space built with the first 3 PCs, which explain 98% of variance. **Inset** shows, as a function of *d*_1_, the Euclidean distances between projections, *P*_*j*_, (where *j* = 1, …, 5 labels the possible values of *d*_1_ in the diagonal block) taken as reference the projection of the shortest *d*_1_. The compression of this curved manifold (see definition in Methods, Eq. 52) was *C*_25_[*P*] = 0.57 while the noiseless case was *C*_0_[*P*] = 0.53 (Fig. 7a, Inset). These values were significantly different (p-value = 1.2e-07; Mann-Wilcoxon test over two bootstrap distributions, each with 100 resamples). **b**. Linear regression of the projections (*aP*_*j*_ + *b*) to the true *d*_1_ (left), the Bayesian estimator (middle) and the ML estimator (right), as a function of *d*_1_. Colored dots show multiple estimates of the regression derived from bootstrapping (*n*=100). Solid lines indicate the true *d*_1_ (left), the Bayesian estimator (middle) and the ML estimator (right) versus *d*_1_. **c**. Violin plots (kernel probability density) on shufflings of the RMSE differences between the three hypotheses (*d*_1_, Bayesian estimator and ML estimator). The width of the gray area indicates the density of the shufflings located there. Horizontal line inside of each violin plot indicates the distribution mean. **d-f**. Mean relative distances during the delay period, which activity is corrupted with Gaussian noise of SD=25 mV. **d**. Relative distance, *D*_*j*_(*t*), between each trajectory *j* = 1, …, 5 and the reference trajectory *d*_1_=370 ms. Neural trajectories are embedded in a state space built with 3 PCs, which explain 91% of variance. Shadings indicate the mean ± 95% CIs computed from 100 bootstrap resamples. **Inset** shows the temporal means of relative distances as a function of *d*_1_. The associated compression, *C*_25_[*D*] = 0.46, was significantly different from that of the noiseless case (Fig. 7 d, Inset), *C*_0_[*D*] = 0.26 (p-value = 7.6e-33). **e**. Linear regression of the mean relative distances (*aD*_*j*_ + *b*) to the true *d*_1_ (left), the Bayesian estimator (middle) and the ML estimator (right) as a function of *d*_1_ (same format as in (b)). **f**. Analogous to (c), but for the relative distance analysis.

**Extended Data Fig. 9.**
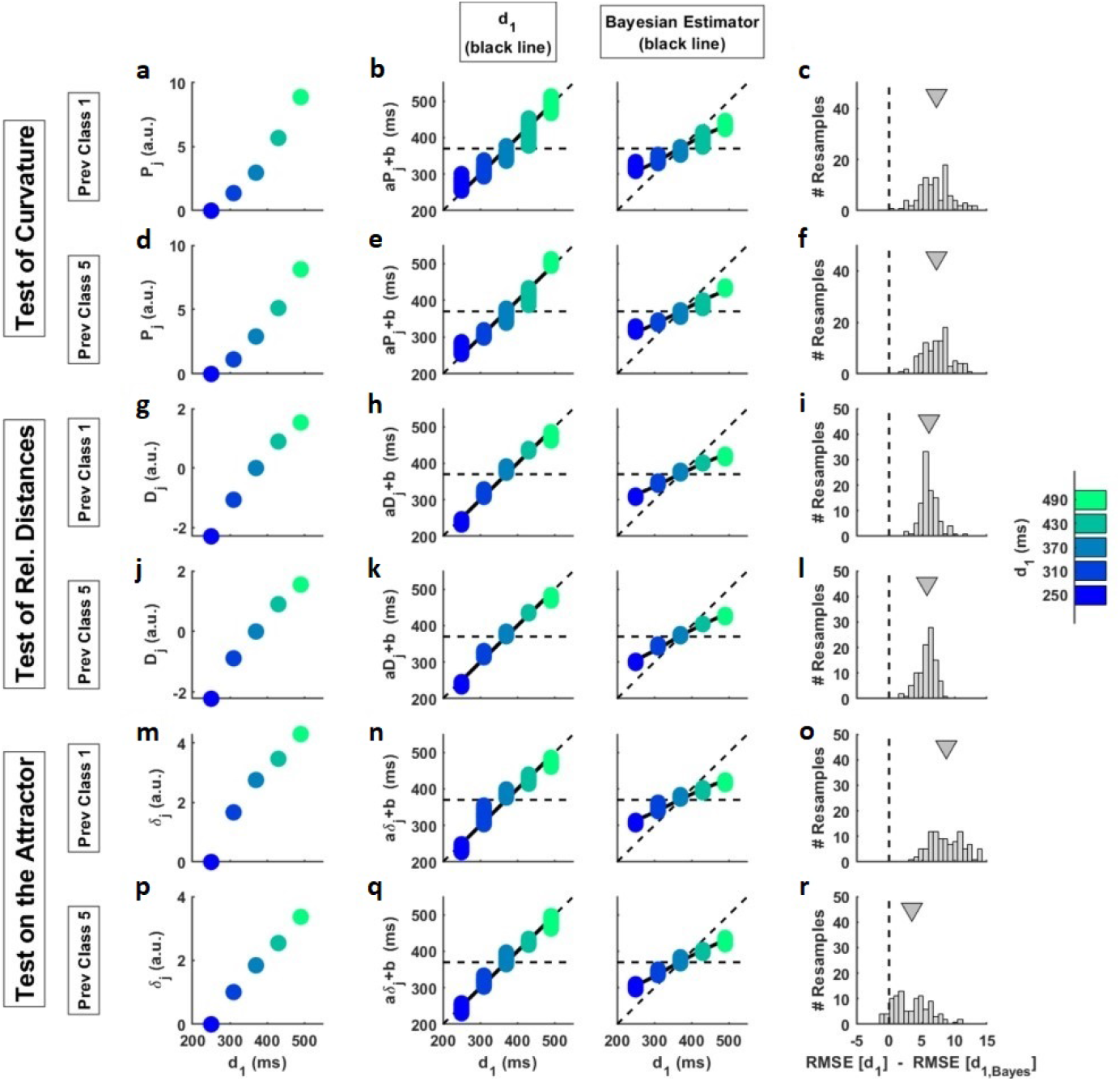
Effect of stimulation history on state-space geometry and comparison with behavior. **a-r**. Example network 1. **a-c**. State-space geometry and test during the presentation of the first interval and conditioning over the previous class 1. **a**. Euclidean distances between projections, *P*_*j*_, (where *j* = 1, …, 5 labels the possible values of *d*_1_ in the diagonal block) of each SF1 state onto the encoding axis, **u**, as a function of *d*_1_. **b**. Fit of a linear regression of the distances between projections (*aP*_*j*_ + *b*) to the true *d*_1_ (left) and the Bayesian estimator (right) as a function of *d*_1_. Colored circles show multiple estimates of the regression derived from bootstrapping (*n*=100). Solid lines indicate the true *d*_1_ (left) and the Bayesian estimator (right) as a function of *d*_1_. **c**. Histogram on shufflings of the difference of the RMSEs between the true *d*_1_ hypothesis and the Bayesian estimator hypothesis. Gray triangles indicate the mean of each histogram. Dashed line identifies 0. **d-f**. State-space geometry and test during the presentation of the first interval and conditioning over the previous class 5 (same format as in (a-c)). **g-l**. State-space geometry and test using the temporal mean of the relative distances, *D*_*j*_, during the delay-period and conditioning over the previous class 1 (**g-i**) and over the previous class 5 (**j-l**). Same format as in (a-c). **m-r**. State-space geometry and test using the distances, *δ*_*j*_, between consecutive terminal points of the trajectories on the attractor and conditioning over the previous class 1 (**m-o**) and over the previous class 5 (**p-r**). Same format as in (a-c). The tests over curvature and relative distances were done with 5-9 PCs to explain at least 90% of the variance. The attractor tests only required the first 2 PCs to explain 99% of the variance.

