## Supplementary Information for "Emergent perceptual biases from state-space geometry in spiking recurrent neural networks trained to discriminate time intervals"

**Luis Serrano-Fernandez, Manuel Beiran, Nestor Parga**

### Supplementary Figures

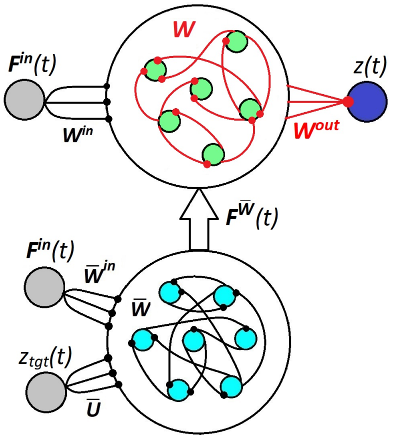


**Supplementary Fig. S1. Full-FORCE scheme**. The task-performing network (top) is a RNN of $N$ recurrently connected spiking neurons (green circles). The network receives an external input, $\boldsymbol{F^{in}}(t)$, from an input layer (gray circle) connected to the network through the $\boldsymbol{W^{in}}$ synapses. This RNN is also connected to an output layer (dark blue circle) through the couplings $\boldsymbol{W^{out}}$ and generates the output signal $z(t)$. Connections marked in black ($\boldsymbol{W^{in}}$) are non-plastic synapses chosen randomly while the hidden weights $\boldsymbol{W}$ and the output weights $\boldsymbol{W^{out}}$ (both indicated with red lines) are plastic. The target-generating network (bottom) is a network of $\bar{N}$ firing-rate neurons (cyan circles), recurrently connected through non-plastic synapses, $\bar{\boldsymbol{W}}$. This network generates the auxiliary function, $\boldsymbol{F}^{\bar{\boldsymbol{W}}}(t)$, needed as a target to train the intrinsic parameters $\boldsymbol{W}$ of the spiking RNN. It receives as inputs both $\boldsymbol{F^{in}}(t)$ and $z_{tgt}(t)$ through the non-plastic synapses $\bar{\boldsymbol{W}}^{\boldsymbol{in}}$ and $\bar{\boldsymbol{U}}$, respectively. Small circles indicate synapses on post-synaptic neurons.

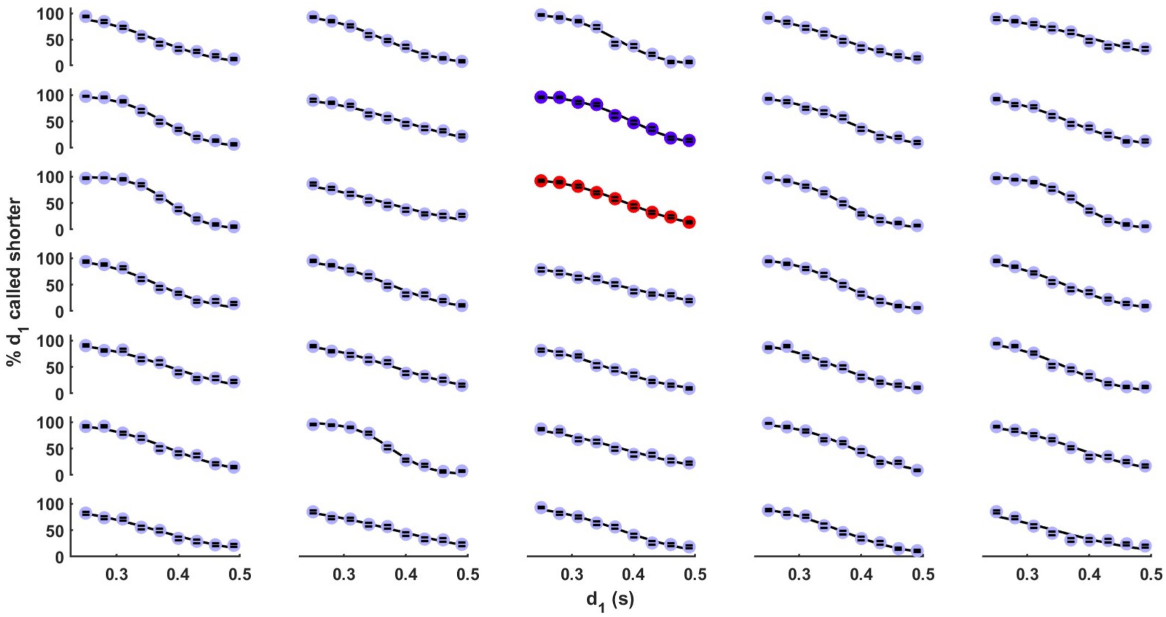


**Supplementary Fig. S2. Psychometric curves are heterogeneous across the 35 trained networks**. Psychometric curves were computed from the horizontal classes of the full block and error bars were obtained from 1000 bootstrap resamples. Black lines indicate sigmoidal fits. Example network 1 (2) appears in the panel with dark blue (red) circles.

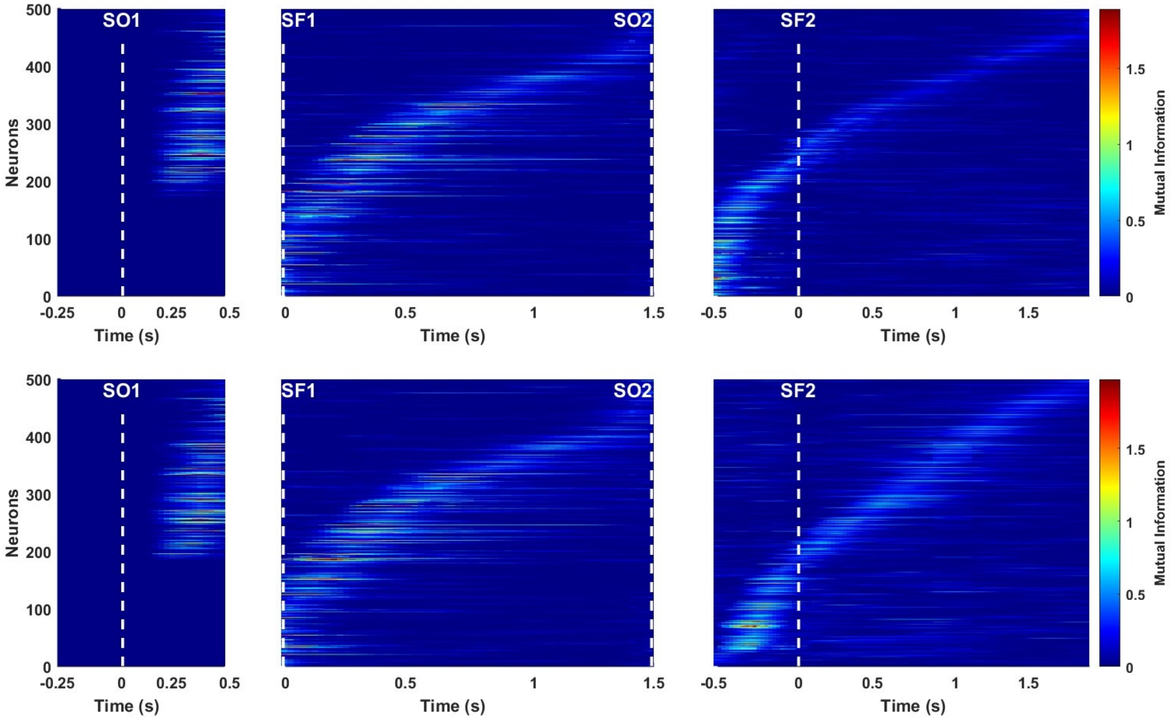


**Supplementary Fig. S3. Mutual Information (MI) about the current $d_1$ of the $N$=500 cells from example network 1. Upper.** MI as a function of time from rewarded trials with choice "$d_1 < d_2$". **Bottom**. MI as a function of time from rewarded trials with choice "$d_1 > d_2$". Three aligments are presented: at the onset of the first stimulus (SO1. Left); at the offset of the first stimulus (SF1. Middle) and at the onset of the second stimulus (SO2. Right). Non-significant MI values (see Methods) are represented in dark blue.

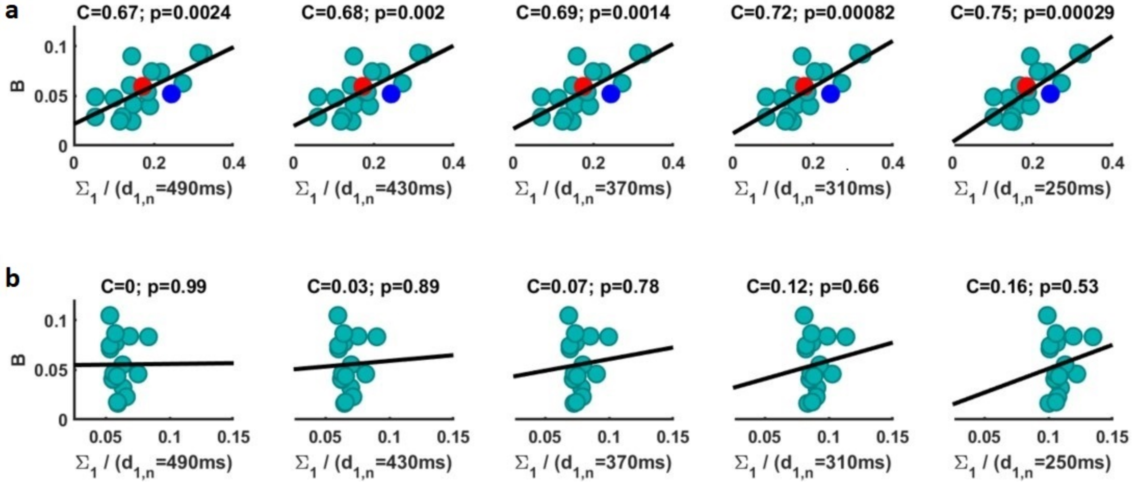


**Supplementary Fig. S4. Correlations between phenomenological bias ($B$) and the total variance of observation probability ($\Sigma_1$).** Correlations are given for all values of the first stimulus of the current ($n$) trial. **a**. Correlations are over the networks classified as positive by model version $m = 0$. Blue circles indicate example network 1. Red circles refer to example network 2. **b**. Correlations are over the networks classified as negative by model version $m = 0$.

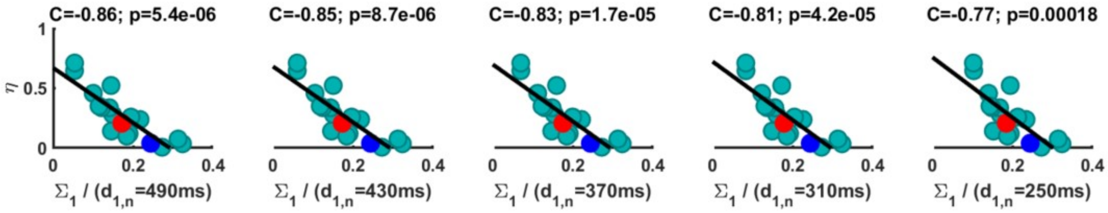


**Supplementary Fig. S5. Correlations between the sensory history parameter ($\eta$) and the total variance of observation probability ($\Sigma_1$).** Correlations are given for all values of the first stimulus of the current ($n$) trial. Correlations are over the networks classified as positive by model version $m = 0$. Blue circles indicate example network 1. Red circles refer to example network 2.

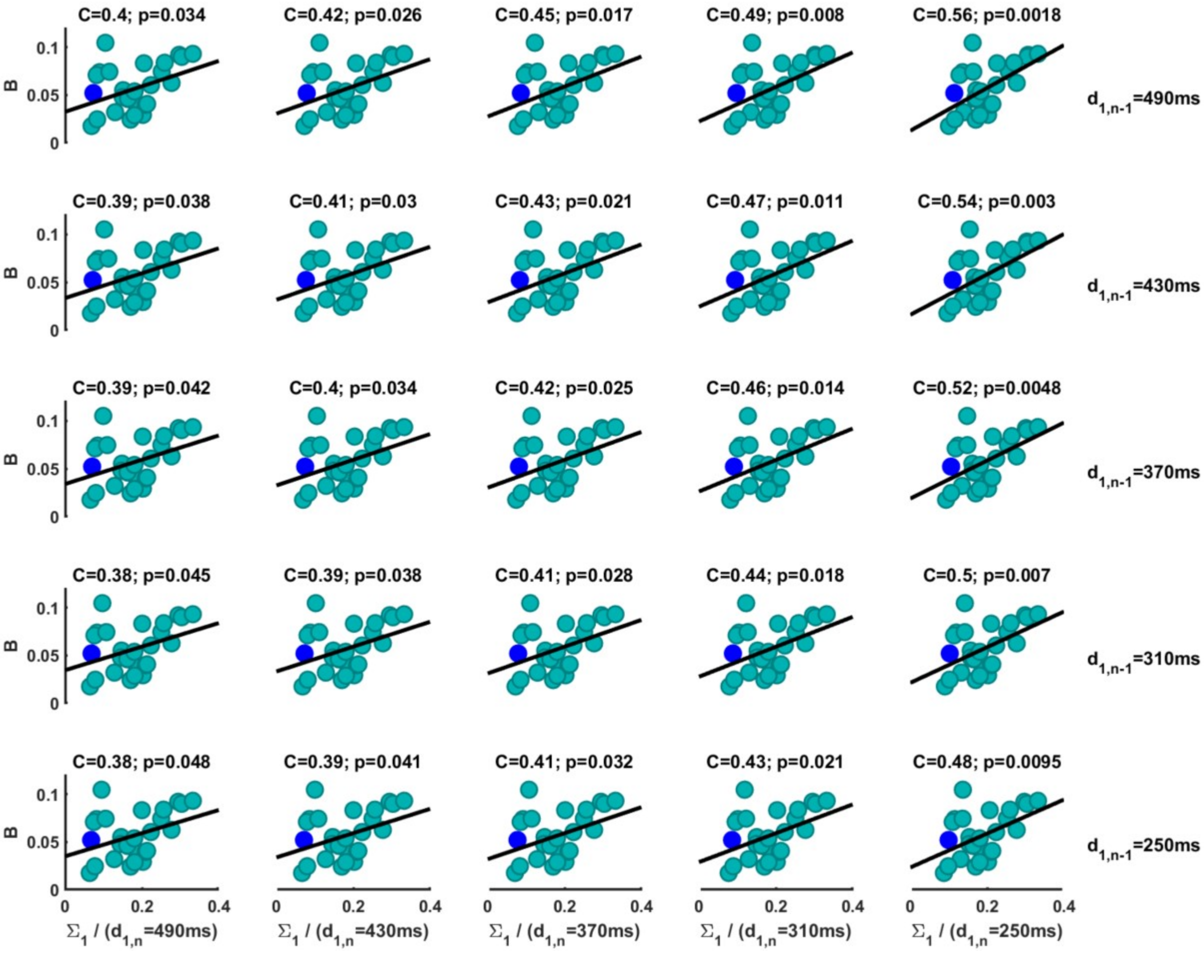


**Supplementary Fig. S6. Correlations between the phenomenological bias ($B$) and the total variance of observation probability ($\Sigma_1$).** Correlations are given for all values of the first stimulus of the current ($n$) and previous ($n-1$) trials. Correlations are over the networks classified as positive by model version $m = 1$. Blue circles indicate example network 1.

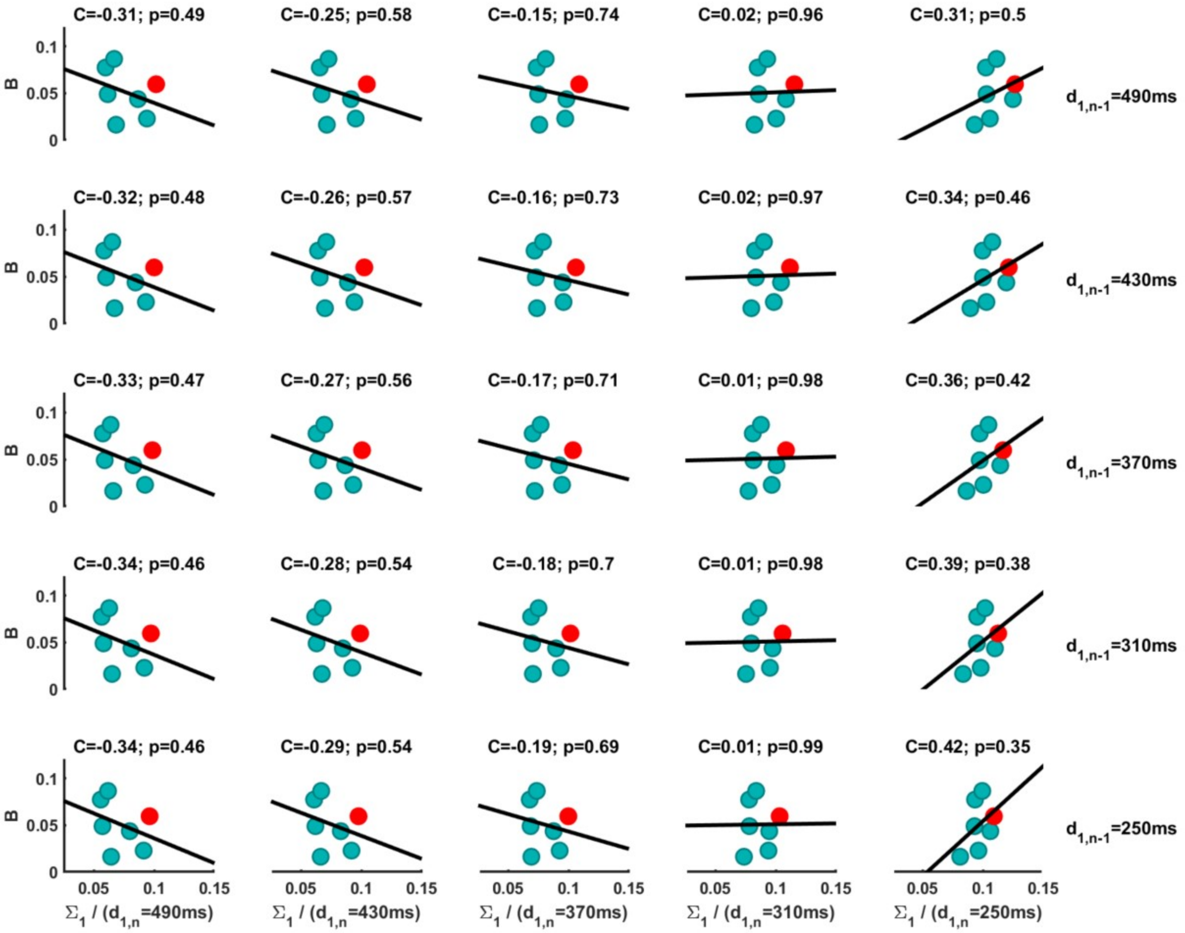


**Supplementary Fig. S7. Correlations between the phenomenological bias ($B$) and the total variance of observation probability ($\Sigma_1$).** Correlations are given for all values of the first stimulus of the current ($n$) and previous ($n-1$) trials. Correlations are over the networks classified as negative by model version $m = 1$. Red circles indicate example network 2.

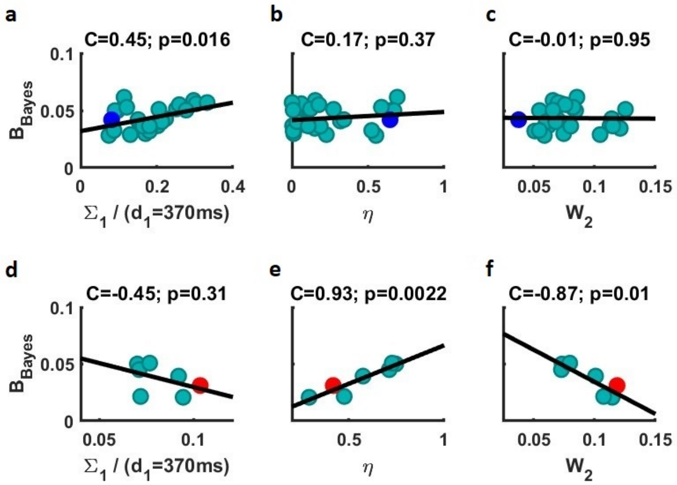


**Supplementary Fig. S8. Correlations between the bias of $d_{1,Bayes}$ ($B_{Bayes}$) and the model parameters**. **a-c**. Correlations over the 28 networks classified as positive by model version $m = 1$. Blue circles indicate example network 1. **a**. Correlation between $B_{Bayes}$ and the total variance of observation probability ($\Sigma_1$) for the current and previous first stimulus equal to 370 ms (the rest of the cases hardly change). **b**. Correlation between $B_{Bayes}$ and the sensory history parameter ($\eta$). **c**. Correlation between $B_{Bayes}$ and the noise parameter of $d_2$ measurements ($W_2$). **d-f**. Correlations over the 7 networks classified as negative by model version $m = 1$. Red circles indicate example network 2. **d**. Correlation between $B_{Bayes}$ and $\Sigma_1$ for the current and previous first stimulus equal to 370 ms (the rest of the cases hardly change). **e**. Correlation between $B_{Bayes}$ and $\eta$. **f**. Correlation between $B_{Bayes}$ and $W_2$. C indicates correlation values and p is the p-value.

### Supplementary Tables

| Model | $\mathbf{W_1}$ | $\mathbf{W_2}$ | $\eta$ | $\gamma_1$ | $\gamma_2$ | RMSE |
|---|---|---|---|---|---|---|
| m=1 without $d_{2,n-1}$ | 0.15 | 0.038 | 0.64 | 0.45 | 0.084 | 0.0011 |
| m=1 with $d_{2,n-1}$ | 0.19 | 0.048 | 0.57 | 0.46 | 0.22 | 0.0012 |

**Supplementary Table S1. Model parameters and RMSE values from model version m=1 including and not including the previous second interval,** $d_{2,n-1}$. Fitted behavioral data corresponds to the example network 1. $W_1$ and $W_2$ are the uncertainties in the observations of the first and second stimulus, respectively. $\eta$ is the history parameter. $\gamma_1$ and $\gamma_2$ are the two model parameters that include a win-stay and loss-shift strategy (WSLS) in the decision making (see Methods). RMSE is the root mean square error from the difference between the behavioral data performance and the model performance.

| Net | BIC with $\mu$ | BIC without $\mu$ | $\Delta$BIC | AIC with $\mu$ | AIC without $\mu$ | $\Delta$AIC |
|---|---|---|---|---|---|---|
| 1 | 4224 | 4171 | 53 | 4205 | 4159 | 46 |
| 2 | 1047 | 1039 | 8 | 1028 | 1026 | 2 |
| 3 | 900 | 872 | 28 | 881 | 859 | 22 |
| 4 | 2160 | 2157 | 3 | 2141 | 2145 | -4 |
| 5 | 1523 | 1515 | 8 | 1504 | 1502 | 2 |
| 6 | 6981 | 6969 | 12 | 6963 | 6956 | 7 |
| 7 | 2129 | 2033 | 96 | 2110 | 2021 | 89 |

**Supplementary Table S2. Comparison between the model with knowledge of the interval mean and the model without it**. The weight of this knowledge is given by the parameter $\mu$. Seven networks trained and tested with reset were used. The criteria AIC and BIC (Methods) tend to favor the model that does not require knowledge of the mean of the intervals.

### Supplementary Texts

#### Supplementary Text S1.

*A normative model that includes the second interval.* Here we present a more general model than the one considered in the main text. This model includes, in addition to the sensory history of the first stimulus ($d_1$), the history of the second interval ($d_2$). Thus, we assumed that at the current trial ($n$) the Bayesian observer makes an observation of the first interval ($o_{1,n}$) given by a combination of the current-trial noisy measurement of $d_1$ ($d'_{1,n}$), measurements of the first and second intervals presented in the $m$ previous trials and observations at a longer temporal scale ($o_{1,n-(m+1)}$)

$$o_{1,n} = \sum_{j=0}^{m} \eta^{2j} (1-\eta)\, d'_{1,n-j} + \sum_{j=0}^{m} \eta^{2j+1} (1-\eta)\, d'_{2,n-(1+j)} + \eta^{2(1+m)} o_{1,n-(m+1)}, \tag{S1}$$

where the parameter $\eta$ measures the effect of the contribution from the activity of the previous trials. We assumed that the noisy measurements of the intervals presented in trials $n-j$ ($d_{1,n-j}$, where $j = 0, 1, ..., m$) are described by normal distributions

$$d'_{1,n-j} \sim \mathcal{N}\left(d_{1,n-j}, W_1^2 d_{1,n-j}^2\right), \tag{S2}$$

and same for the second stimulus in trials $n-(1+j)$ ($d_{2,n-(1+j)}$ where $j = 0, 1, ..., m$)

$$d'_{2,n-(1+j)} \sim \mathcal{N}\left(d_{2,n-(1+j)}, W_2^2 d_{2,n-(1+j)}^2\right), \tag{S3}$$

where $W_1$ and $W_2$ are the Weber fractions, two dimensionless parameters of the Bayesian model to be determined. Another assumption is that trials prior to the last $m$ constitute a Gaussian stationary regime (the long-term sensory history). The mean and variance of the stationary variable are obtained using self-consistency

$$M_{12,st} = \frac{\langle d_1 \rangle + \eta \langle d_2 \rangle}{1+\eta} \tag{S4}$$

$$\Sigma_{12,st}^2 = \frac{1-\eta}{1+\eta} \left[ \frac{W_1^2 \langle d_1^2 \rangle + \sigma_{d_1}^2 + \eta^2 \left( W_2^2 \langle d_2^2 \rangle + \sigma_{d_2}^2 \right)}{1+\eta^2} \right], \tag{S5}$$

where $\langle d_2 \rangle$ and $\sigma_{d_2}^2$ are the mean and variance of the prior probability of $d_2$. Then, the observation of $d_1$ in the current trial, described by the variable $\mathcal{O}_1$, is also a Gaussian variable with mean $\mathcal{M}$ and variance $\Sigma_1^2$. Its mean and variance are

$$\mathcal{M} = (1-\eta)\sum_{j=0}^{m}\left[\eta^{2j}d_{1,n-j} + \eta^{2j+1}d_{2,n-(1+j)}\right] + \eta^{2(1+m)}\frac{\langle d_1\rangle + \eta\langle d_2\rangle}{1+\eta} \tag{S6}$$

$$\Sigma_1^2 = (1-\eta)^2\sum_{j=0}^{m}\eta^{4j}\left[W_1^2 d_{1,n-j}^2 + \eta^2 W_2^2 d_{2,n-(1+j)}^2\right] + \eta^{4(1+m)}\Sigma_{12,st}^2. \tag{S7}$$

Notice that if we define the normalized means:

$$d_{12,n} = \frac{d_{1,n} + \eta d_{2,n-1}}{1+\eta}$$

$$\langle d_{12}\rangle = \frac{\langle d_1\rangle + \eta\langle d_2\rangle}{1+\eta}$$

$$d_{12,n-j} = \frac{d_{1,n-j} + \eta d_{2,n-(1+j)}}{1+\eta},$$

then the mean of the likelihood of an observation $\mathcal{O}_1$ can be expressed as

$$\mathcal{M} = \left(1-\eta^2\right)\sum_{j=0}^{m}\left[\eta^{2j}d_{12,n-j}\right] + \eta^{2(1+m)}\langle d_{12}\rangle \tag{S8}$$

$$\mathcal{M} = d_{12,n} + \sum_{j=1}^{m}\eta^{2j}\left[d_{12,n-j} - d_{12,n-(j-1)}\right] + \eta^{2(1+m)}\left[\langle d_{12}\rangle - d_{12,n-m}\right] \tag{S9}$$

Also in the general model the weights of the various contributions to the mean are normalized. Again, this is reminiscent of a result by [1] who found that the weights in their best multilinear regression model appear normalized.

The particular case in which the transient regime contains only the class in the previous trial (i.e., $m = 1$) has the following mean and variance of the observation probability

$$\mathcal{M} = d_{12,n} + \eta^2\left[d_{12,n-1} - d_{12,n} + \eta^2\left(\langle d_{12}\rangle - d_{12,n-1}\right)\right] \tag{S10}$$

$$\Sigma_1^2 = (1-\eta)^2\left[W_1^2 d_{1,n}^2 + \eta^4 W_1^2 d_{1,n-1}^2\right] + \eta^2(1-\eta)^2\left[W_2^2 d_{2,n-1}^2 + \eta^4 W_2^2 d_{2,n-2}^2\right] + \eta^8\Sigma_{12,st}^2. \tag{S11}$$

The likelihood $P\left(\mathcal{O}_1|d_{1,n}, d_{1,n-1}, d_{2,n-1}, d_{2,n-2}\right)$, of an observation $\mathcal{O}_1$ given $d_{1,n}$, $d_{1,n-1}$, $d_{2,n-1}$ and $d_{2,n-2}$, has the mean and variance in Eq. (S10) and Eq. (S11), respectively. Given the prior of the second stimulus, it is trivial to marginalize it over $d_{2,n-2}$, obtaining $P\left(\mathcal{O}_1|d_{1,n}, d_{1,n-1}, d_{2,n-1}\right)$. Now, it is possible to calculate the posterior, $P\left(d_{1,n}, d_{1,n-1}, d_{2,n-1}|\mathcal{O}_1\right) \equiv P(d_{1,n}, C_{n-1}|\mathcal{O}_1)$

$$P\left(d_{1,n}, C_{n-1} | \mathcal{O}_1\right) \propto P\left(\mathcal{O}_1 | d_{1,n}, C_{n-1}\right) P\left(d_{1,n}\right) P\left(C_{n-1}\right), \tag{S12}$$

where $C_{n-1}$ indicates the previous class $(d_{1,n-1}, d_{2,n-1})$. Finally, the noisy observation of $d_2$, denoted as $\mathcal{O}_2$, was taken as a Gaussian variable with mean $d_2$ and standard deviation $\sigma_2 = W_2 d_2$,

$$\mathcal{O}_2 \sim \mathcal{N}\left(d_2, W_2^2 d_2^2\right). \tag{S13}$$

The distribution $P\left(\mathcal{O}_2 | d_{2,n}\right)$ defines the likelihood of an observation $\mathcal{O}_2$, given the current trial second interval.

#### Supplementary Text S2.

*Marginalization over previous class and outcome.* The objective of this Supplementary Text is to obtain the marginal discrimination curve $P(U_0 = 1|C_0^k)$ $(k = 1, \ldots, 10)$ by marginalizing $P(S|C_0^k,\, C_{-1}^l,\, U_{-1} = 1)$ on the previous class, $C_{-1}^l$, and outcome, $U_{-1}$. We first notice that the probability of deciding $S \equiv$ "$d_1 < d_2$" for given current and previous classes can be decomposed as

$$\begin{aligned} P(S|C_0^k,\, C_{-1}^l) = P(S|C_0^k,\, C_{-1}^l,\, U_{-1} = 1) \cdot P(U_{-1} = 1|C_{-1}^l) + \\ + P(S|C_0^k,\, C_{-1}^l,\, U_{-1} = 0) \cdot P(U_{-1} = 0|C_{-1}^l). \end{aligned} \tag{S14}$$

Marginalizing on the previous class $l$

$$\begin{aligned} \sum_{l=1}^{10} \frac{1}{10} P(S|C_0^k) = \sum_{l=1}^{10} \frac{1}{10} \Big[ P(S|C_0^k,\, C_{-1}^l,\, U_{-1} = 1) \cdot P(U_{-1} = 1|C_{-1}^l) + \\ + P(S|C_0^k,\, C_{-1}^l,\, U_{-1} = 0) \cdot P(U_{-1} = 0|C_{-1}^l) \Big]. \end{aligned} \tag{S15}$$

We will first deal with the case in which the current class $k$ is on the upper diagonal $(k = 1, \ldots, 5)$. Then, using that $P(S|C_0^k) = P(U_0 = 1|C_0^k)$ we obtain

$$\begin{aligned} 10 P(U_0 = 1|C_0^k) = \sum_{l=1}^{10} \Big[ P(S|C_0^k,\, C_{-1}^l,\, U_{-1} = 1) \cdot P(U_{-1} = 1|C_{-1}^l) + \\ + P(S|C_0^k,\, C_{-1}^l,\, U_{-1} = 0) \cdot P(U_{-1} = 0|C_{-1}^l) \Big]. \end{aligned} \tag{S16}$$

Moreover, using that $P(U_{-1} = 0|C_{-1}^l) = 1 - P(U_{-1} = 1|C_{-1}^l)$, we have

$$10P(U_0=1|C_0^k) = \sum_{l=1}^{10}\Big[P(S|C_0^k,\, C_{-1}^l,\, U_{-1}=1)\cdot P(U_{-1}=1|C_{-1}^l) + \\ + P(S|C_0^k,\, C_{-1}^l,\, U_{-1}=0)\cdot\Big(1-P(U_{-1}=1|C_{-1}^l)\Big)\Big]. \tag{S17}$$

Reorganizing this equation we have, for a given current class $k$

$$\sum_{l=1}^{10}\Big[P(S|C_0^k,\, C_{-1}^l,\, U_{-1}=1) - P(S|C_0^k,\, C_{-1}^l,\, U_{-1}=0)\Big]\cdot P(U_{-1}=1|C_{-1}^l) \\ - 10P(U_0=1|C_0^k) = -\sum_{l=1}^{10} P(S|C_0^k,\, C_{-1}^l,\, U_{-1}=0).$$

Using that $P(U_{-1}=1|C_{-1}^l) = P(U_0=1|C_0^l)$, we have

$$\sum_{l=1}^{10}\Big[P(S|C_0^k,\, C_{-1}^l,\, U_{-1}=1) - P(S|C_0^k,\, C_{-1}^l,\, U_{-1}=0)\Big]\cdot\, P(U_0=1|C_0^l) \\ - 10P(U_0=1|C_0^k) = -\sum_{l=1}^{10} P(S|C_0^k,\, C_{-1}^l,\, U_{-1}=0). \tag{S18}$$

These are 5 equations ($k=1,\ldots,5$) with 10 unknowns $P(U_0=1|C_0^l)$ ($l=1,\ldots,10$). On the other hand, for current classes $k$ on the bottom diagonal ($k=6,\ldots,10$), we obtain the following 5 equations, one for each current class located on the bottom diagonal

$$\sum_{l=1}^{10}\Big[P(S|C_0^k,\, C_{-1}^l,\, U_{-1}=1) - P(S|C_0^k,\, C_{-1}^l,\, U_{-1}=0)\Big]\cdot\, P(U_0=1|C_0^l) \\ + 10P(U_0=1|C_0^k) = 10 - \sum_{l=1}^{10} P(S|C_0^k,\, C_{-1}^l,\, U_{-1}=0). \tag{S19}$$

Solving the system in Eqs. (S18-S19) we obtain the marginal accuracy curve, $P(U_0=1|C_0^l)$ ($l=1,\ldots,10$) (Fig. 5g). Once this is known, using Eq. (S14) it is possible to compute the probability of deciding $S \equiv d_1 < d_2$" conditioned on the current and the previous classes $P(S|C_0,\, C_{-1})$, and from it to obtain the accuracy curves in Fig. 5d.
